# Mitochondrial nucleoid organization and biogenesis of complex I require mTERF18/SHOT1 and ATAD3 in *Arabidopsis thaliana*

**DOI:** 10.1101/2020.05.11.088575

**Authors:** Minsoo Kim, Vincent Schulz, Lea Brings, Theresa Schoeller, Kristina Kühn, Elizabeth Vierling

**Author notes:** Corresponding Author: Elizabeth Vierling, Department of Biochemistry and Molecular Biology, University of Massachusetts, Amherst, MA 01003.

## Abstract

Mitochondria play critical roles in eukaryotes in ATP generation through oxidative phosphorylation (OXPHOS) and also produce both damaging and signaling reactive oxygen species (ROS). Originating from endosymbiosis, mitochondria have their own reduced genomes that encode essential subunits of the OXPHOS machinery. MTERF (Mitochondrial Transcription tERmination Factor-related) proteins have been shown to be involved in organelle gene expression by interacting with organellar DNA or RNA in multicellular eukaryotes. We previously identified mutations in *Arabidopsis thaliana MTERF18/SHOT1* that enable plants to better tolerate heat and oxidative stresses, presumably due to low ROS and reduced oxidative damage. To understand molecular mechanisms leading to *shot1* phenotypes, we investigated mitochondrial defects of *shot1* mutants and targets of the SHOT1 protein. *shot1* mutants have problems accumulating OXPHOS complexes that contain mitochondria-encoded subunits, with complex I and complex IV most affected. SHOT1 binds specific mitochondrial DNA sequences and localizes to mitochondrial nucleoids, which are diffuse in *shot1* mutants. Furthermore, three homologues of mammalian ATAD3A proteins, which are suggested to be involved in mitochondrial nucleoid organization, were identified as SHOT1-interacting proteins (designated SHOT1 BINDING ATPASES (SBA)1, 2 and 3). Importantly, disrupting SBA function also disrupts nucleoids, compromises accumulation of complex I and enhances heat tolerance. We conclude that proper nucleoid organization is critical for correct expression and accumulation of complex I, and propose that nucleoid disruption results in unique changes in mitochondrial metabolism and signaling that lead to heat tolerance.

**Significance:** In all eukaryotes, mitochondria are critical organelles that supply chemical energy for life, which is produced by the oxidative phosphorylation (OXPHOS) machinery on the inner mitochondrial membrane. The OXPHOS machinery comprises multiple protein complexes with subunits encoded by both nuclear and mitochondrial genes. Nuclear-encoded mTERF proteins are important for expression of mitochondrial genes, interacting with mitochondrial DNA or RNA. Our study reveals that the Arabidopsis mTERF18/SHOT1 protein interacts with mtDNA and homologs of human ATAD3A proteins, and that both proteins are critical for mitochondrial nucleoid organization and accumulation of OXPHOS Complex I. Further, the data indicate nucleoid disruption leads to unique mitochondrial and cellular responses such that mutant plants have enhanced heat tolerance.

## Introduction

As a metabolic hub in eukaryotic cells, mitochondria generate energy in the form of ATP by oxidative phosphorylation (OXPHOS), as well as produce key metabolites necessary for cellular homeostasis and growth. Mitochondria are also the source of both signaling and damaging reactive oxygen species (ROS), such that control of mitochondrial metabolism is critical to life. After taking up residence in ancestral eukaryotic cells, mitochondria lost most of their DNA over billions of years of evolution. The small number of genes retained in mitochondria encode subunits of OXPHOS complexes I, III, IV and V, rRNAs and some tRNAs (1, 2). Additionally, mitochondrial DNA (mtDNA) in plants contains genes encoding some mitochondrial ribosomal proteins and cytochrome *c* maturation factors. Most of the proteins (∼2000) required for mitochondrial function are supplied by nuclear genes (3). Consequently, mitochondria require their own gene expression system that must be coordinated with nuclear gene expression for biogenesis of the OXPHOS machinery.

In plants, but not in animals, mitochondrial pre-mRNAs undergo complex post-transcriptional processes that include RNA trimming, processing of polycistronic transcripts, cytidine to uridine editing and splicing (4). These processes require hundreds of nuclear-encoded proteins. Eukaryotic protein families that contribute to these processes have particularly expanded in plants, including the mTERF (mitochondrial transcription termination factor) proteins (5). mTERFs, which contain variable numbers of a ∼30 amino acid motif (6), were first discovered in human mitochondria and are unique to multicellular eukaryotes (7). All mTERFs studied to date bind either DNA or RNA in organelles (both mitochondria and plastids in plants) and control DNA replication or organelle gene expression at different levels (4, 8–10). In *Arabidopsis thaliana*, there are 35 mTERFs, most of which are localized to chloroplasts and/or mitochondria (5, 11). Loss-of-function mutations in mTERF1/SINGLET OXYGEN-LINKED DEATH ACTIVATOR10 (SOLDAT10) and mTERF4/BELAYA SMERT (BSM)/RUGOSA2 (RUG2) cause arrest in embryo development (11–13), and knock-out of mTERF6 causes seedling lethality (14), highlighting mTERF importance in plant development. Furthermore, chloroplast-localized mTERFs, such as mTERF5/MDA1 and mTERF9/TWIRT1, are essential for normal chloroplast development, as well as being involved in abiotic stress tolerance (15, 16).

There is limited information on the molecular functions of plant mTERFs. Maize mTERF4 participates in splicing chloroplast group II introns, which is probably conserved in *A. thaliana* mTERF4/BSM/RUG2 (11, 17). MTERF15 is involved in splicing mitochondrial *nad2* intron 3, and mutants show retarded growth (18). mTERF6 binds chloroplast isoleucine tRNA (*trnI.2*) during its maturation (14), and binds DNA at the 3’ region of the *rpoA* polycistron, likely acting as a transcription terminator (19). MTERF8/pTAC15 (plastid Transcriptionally Active Chromosome protein) is also involved in transcription termination, binding the 3’-end of chloroplast *psbJ* (20). mTERF5 positively regulates chloroplast *psbEFLJ* transcription (21), and MTERF22 affects mitochondrial transcription by an unidentified mechanism (22). For most plant mTERFs their nucleic acid targets and interacting proteins remain to be identified.

We previously isolated a mutant of *A. thaliana mTERF18/SHOT1* (*Suppressor #1 of HOT1-4;* AT3G60400) in a suppressor screen of a heat sensitive mutant of the molecular chaperone HSP101(AT1G74310) (23). *shot1* mutants exhibit retarded growth presumably due to mitochondrial dysfunction, but increased themotolerance likely resulting from reduced ROS and reduced oxidative damage during heat stress. Here, we show that SHOT1 is required for normal accumulation of OXPHOS complexes I, III, IV and V, all of which contain at least one mitochondria-encoded subunit. We demonstrate SHOT1 is associated with mtDNA, localizes to mitochondrial nucleoids and is required for proper nucleoid organization. Furthermore, SHOT1 interacts with three homologues of mammalian ATAD3A proteins, which are mitochondrial membrane ATPases reported to interact with nucleoids; altering their function mimics a number of *shot1* phenotypes. We conclude that proper mitochondrial nucleoid organization is critical for OXPHOS accumulation, mitochondrial morphology and function.

## Results

### *shot1* mutants are impaired in OXPHOS biogenesis

As an mTERF protein, SHOT1 is expected to be involved in mitochondrial gene expression or mtDNA maintenance. Previously, to find possible SHOT1 targets, we determined mitochondrial transcript levels in a recessive *shot1* mutant in *A. thaliana* (23). No specific transcripts were detrimentally affected, indicating that SHOT1 was not directly involved in regulating transcript levels of specific genes. Thus, we initiated analyses of other defects in mitochondria of *shot1* mutants to obtain insight into SHOT1 function. Two recessive mutants of *SHOT1* (*shot1-1* (G105D) and *shot1-2* (T-DNA insertion)) showed retarded growth (Fig. 1A) (23), suggesting a major mitochondrial defect. Therefore, we compared the abundance and activity of OXPHOS complexes between *shot1* mutants and wild type (WT). *shot1* mutants show a reduction of complexes I and V (Fig. 1B, left), and a corresponding decrease in complex I activity (Fig. 1B, middle). In addition, cytochrome oxidase (complex IV) activity is strongly reduced in the mutants (Fig. 1B, right).

**Fig. 1.**
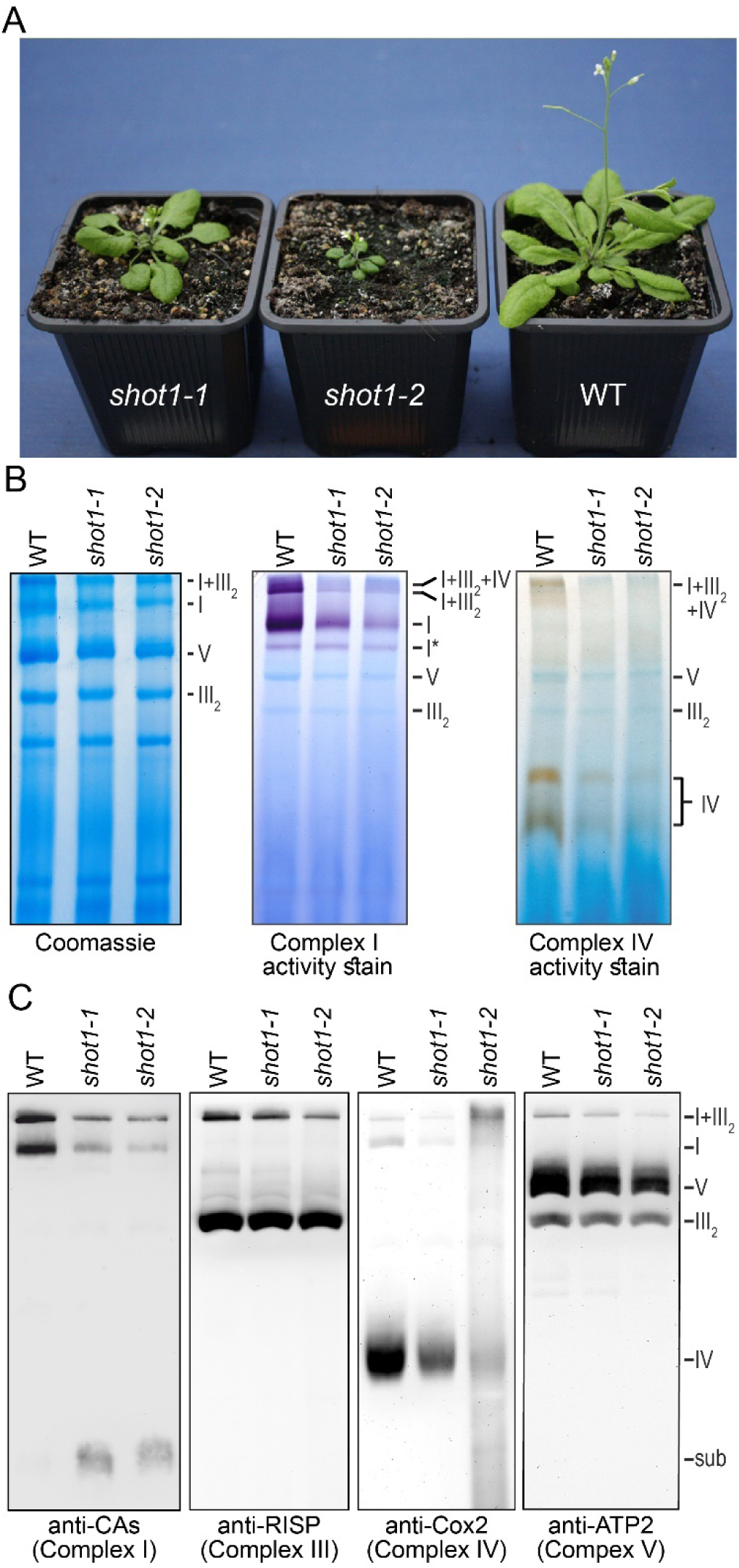
Impaired OXPHOS in *shot1* mutants. (A) 6 week-old *shot1-1, shot1-2* and wild-type plants (Col-0). (B) BN-PAGE analysis of mitochondria, stained with Coomassie or for NADH oxidase or cytochrome oxidase activity. OXPHOS complexes I, III_2_, IV and V and supercomplexes I+III_2_, and I+III_2_+IV are labelled; I* is a complex I assembly intermediate. (C) Immunodetection of BN-PAGE-resolved OXPHOS complexes with indicated antisera: CAs: nucleus-encoded carbonic anhydrase subunits. sub: a likely early assembly intermediate of complex I; RISP: nucleus-encoded Rieske iron-sulfur protein; Cox2: mitochondrion-encoded cytochrome oxidase 2; ATP2: nucleus-encoded beta subunit of ATP synthase. The anti-RISP blot was stripped and probed with anti-ATP2.

Significant reductions in complexes I, IV, V and the I+III_2_ supercomplex and a slight reduction in complex III levels were confirmed by immunoblotting (Fig. 1C and *SI Appendix*, Fig. S1). Accumulation of a complex I assembly intermediate in *shot1* (“sub” in Fig. 1C) supports that reduction of this complex in *shot1* results from impaired biogenesis, rather than decreased complex I stability. This intermediate over-accumulates in plants with reduced expression of the mitochondria-encoded Nad2 subunit (*SI Appendix*, Fig. S2) (24). Its over-accumulation in *shot1* indicates complex I assembly is defective at the level of Nad2 insertion. We conclude that OXPHOS defects in *shot1* are due to either inability to correctly express several mitochondrial genes for OXPHOS subunits, or failure to assemble these subunits into OXPHOS complexes.

### SHOT1 function affects OXPHOS assembly post-translationally

We subsequently examined mitochondrial gene expression in *shot1* mutants in detail, focusing on subunit Nad2 of complex I and the three mitochondria-encoded subunits of complex IV, Cox1, Cox2, and Cox3. RNA gel blots confirmed elevated *nad2, cox1, cox2* and *cox3* transcripts as reported previously (23) and showed WT-like transcript sizes in *shot1*, indicating that synthesis, splicing and end-maturation of these transcripts does not require SHOT1 (*SI Appendix*, Fig. S3). Sequencing cDNAs of *nad2, cox1, cox2*, and *cox3* transcripts showed that *shot1-2* mutants were not defective in editing (*SI Appendix*, Fig. S4). We conclude SHOT1 is not involved in synthesis, splicing, editing or end-maturation of these mitochondrial mRNAs. We then addressed a possible SHOT1 role in translation by examining association of these mRNAs with polysomes (Fig. 2). Polysomal fractions from *shot1-2* seedlings contained at least as much *nad2* and *cox* transcripts as corresponding fractions from WT, and polysomal peaks were not shifted towards lighter fractions in *shot1*. Thus, translation of *nad2* and *cox* transcripts is not impaired in *shot1-2*. Rather, *shot1-2* plants have more mitochondrial ribosomes and polysomes than WT, as indicated by analysis of mitochondrial ribosomal RNAs (Fig. 2), pointing to an overall increase of mitochondrial translation in *shot1-2*.

**Fig. 2.**
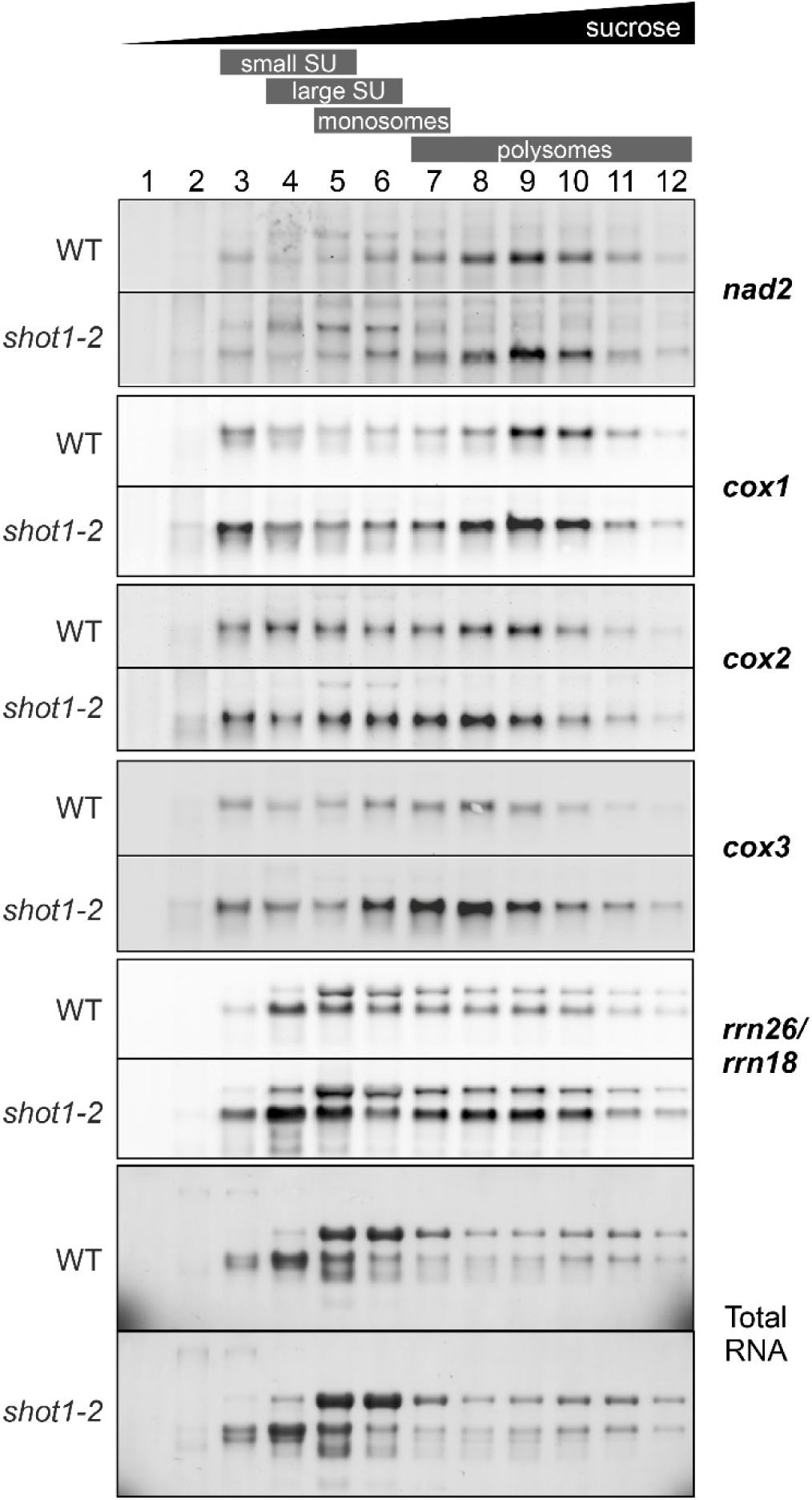
Association of mitochondrial *cox* and *nad2* transcripts with polysomes is unimpaired in *shot1*. Polysomes from total extracts of *shot1-2* and wild-type seedlings were size-fractionated in sucrose gradients and fractions probed by RNA gel blot hybridization. Mitochondrial rRNA probes, *rrn18* and *rrn26*, were hybridized to the same fractions. Total RNA stained with methylene blue is the loading control. The increase in *cox1, cox2, cox3, rrn18* and *rrn26* signals in *shot1-2* compared with the WT represents an increase in mitochondrial ribosomes and polysomes, mitochondrial transcripts, and translated mitochondrial transcripts in the mutant.

Increased mitochondrial transcript levels and elevated protein synthesis in respiratory mutants have been previously observed, e.g., for the *rug3* mutant impaired in complex I biogenesis (25) or the *rpoTmp-1* mutant impaired in both complexes I and IV (26), and are likely secondary effects. We reasoned that SHOT1 does not impact OXPHOS biogenesis through a role in gene expression; rather, its function is important during post-translational steps. We did not consider a potential role for SHOT1 in the expression of TatC, which is the only mitochondrially encoded factor acting in OXPHOS assembly. Mitochondrial TAT system inhibition was recently shown to cause a specific defect in complex III assembly (27) that is not exhibited by *shot1*. Notably, Nad2 is the first mitochondria-encoded protein to enter the complex I assembly pathway (24). Failure in *shot1* mitochondria to perform this complex I assembly step might not necessarily be specific to Nad2 incorporation; it could also reflect a general defect in assembling mitochondria-encoded complex I subunits.

### Mitochondrial proteomic data indicate *shot1 has* major defects in assembly of OXPHOS complexes

Mitochondria contain ∼2,000 proteins, only a small fraction (∼30) of which are encoded in the mitochondrial genome (3, 28). To examine proteome changes resulting from defective SHOT1 and to identify possible SHOT1 targets, we employed label-free quantitative proteomics on isolated mitochondria from *shot1-2* and wild type. A total of 1998 protein groups were detected from all samples. Most proteins related to translation processes were increased in *shot1-2* (Fig. 3A), consistent with the observed increased polysomal mitochondrial RNAs (Fig. 2). Of 33 proteins encoded in the *A. thaliana* mitochondrial genome, 23 were detected (Table 1). Interestingly, in *shot1-2* all mitochondria-encoded subunits of complexes I (seven subunits), III (one subunit), IV (two subunits) and V (four subunits) were significantly decreased, while subunits of cytochrome maturation factor F and ribosomal proteins were increased. Most nucleus-encoded subunits of OXPHOS complexes I, III, IV and V were also decreased (Fig. 3B). Complex I and IV subunits were the most reduced, while there was less decrease in complex V subunits, consistent with BN-PAGE analysis (Fig. 1). Complex III subunits showed a minor decrease, and there were no differences in subunits of complex II, all of which are nucleus-encoded (Fig. 3B and 3C). These results suggest that electron flow through the cytochrome-dependent electron transport chain (ETC), involving complexes I, II, III and IV, would be reduced in the mutant. In plant mitochondria, an alternative ETC pathway, comprising type II NAD(P)H dehydrogenases and an alternative oxidase (AOX), allows electrons to bypass the cytochrome-dependent ETC. As expected, alternative ETC proteins were highly abundant in *shot1-2*, indicating that the mutant relies on alternative ETC (Fig. 3C and S1). The fact that OXPHOS proteins accumulate less in *shot1* mutants than in WT, even though they are translated at a higher level (Fig. 2 and 3A), further strengthens the idea that *shot1* mutants are defective in OXPHOS complex assembly. OXPHOS subunits are known to be degraded when not properly assembled (24) and in accordance with these observations, molecular chaperones and proteases are highly upregulated in *shot1-2* (*SI Appendix*, Fig. S5A).

**Table 1.** Mitochondrion-encoded proteins identified by proteomics. Log2 fold change (log2FC) of *shot1-2* compared to WT with -log10p values are shown.

|  | Identified Proteins (23) | Uniprot ID | log2FC | -log10p |
| --- | --- | --- | --- | --- |
| Complex I | NADH dehydrogenase subunit 1, NAD1 | P92558 | -1.94 | 7.41 |
|  | NADH dehydrogenase subunit 2, NAD2 | O05000 | -2.09 | 7.12 |
|  | NADH dehydrogenase subunit 3, NAD3 | P92533 | -2.04 | 7.84 |
|  | NADH dehydrogenase subunit 4, NAD4 | P93313 | -2.13 | 8.18 |
|  | NADH dehydrogenase subunit 5, NAD5 | P29388 | -2.09 | 7.20 |
|  | NADH dehydrogenase subunit 7, NAD7 | P93306 | -1.18 | 5.35 |
|  | NADH dehydrogenase subunit 9, NAD9 | Q95748 | -0.99 | 5.01 |
| Complex III | Cytochrome b, COB | P42792 | -0.87 | 3.04 |
| Complex IV | Cytochrome c oxidase subunit 1, COX1 | P60620 | -2.10 | 2.50 |
|  | Cytochrome c oxidase subunit 2, COX2 | P93285 | -1.69 | 5.99 |
| Complex V | ATP synthase subunit a | P92547 | -1.41 | 2.87 |
|  | ATP synthase subunit b, ORF25, MI25 | Q04613 | -1.37 | 3.62 |
|  | ATP synthase subunit alpha, ATPA, ATP1 | P92549 | -1.30 | 7.03 |
|  | ATP synthase protein YMF19, ATP8, ORFB | P93303 | -1.21 | 5.49 |
| ccmF | Cytochrome c maturation F N1, CCMFN1 | Q9T6H8 | 0.82 | 1.50 |
|  | Cytochrome c maturation F C, CCMFC | P93286 | 0.81 | 0.90 |
| Ribosomal proteins | Ribosomal protein S3, RPS3 | Q95749 | 1.22 | 6.65 |
|  | Ribosomal protein S4, RPS4 | Q31708 | 1.10 | 3.10 |
|  | Ribosomal protein S7, RPS7 | P92557 | 0.95 | 1.88 |
|  | Ribosomal protein S12, RPS12 | P92532 | 2.52 | 1.83 |
|  | 60S ribosomal protein L2, RPL2 | P93311 | 0.73 | 3.69 |
|  | 60S ribosomal protein L5, RPL5 | P42793 | 1.14 | 3.28 |
|  | 60S ribosomal protein L16, RPL16 | Q95747 | 0.98 | 1.70 |

**Fig. 3.**
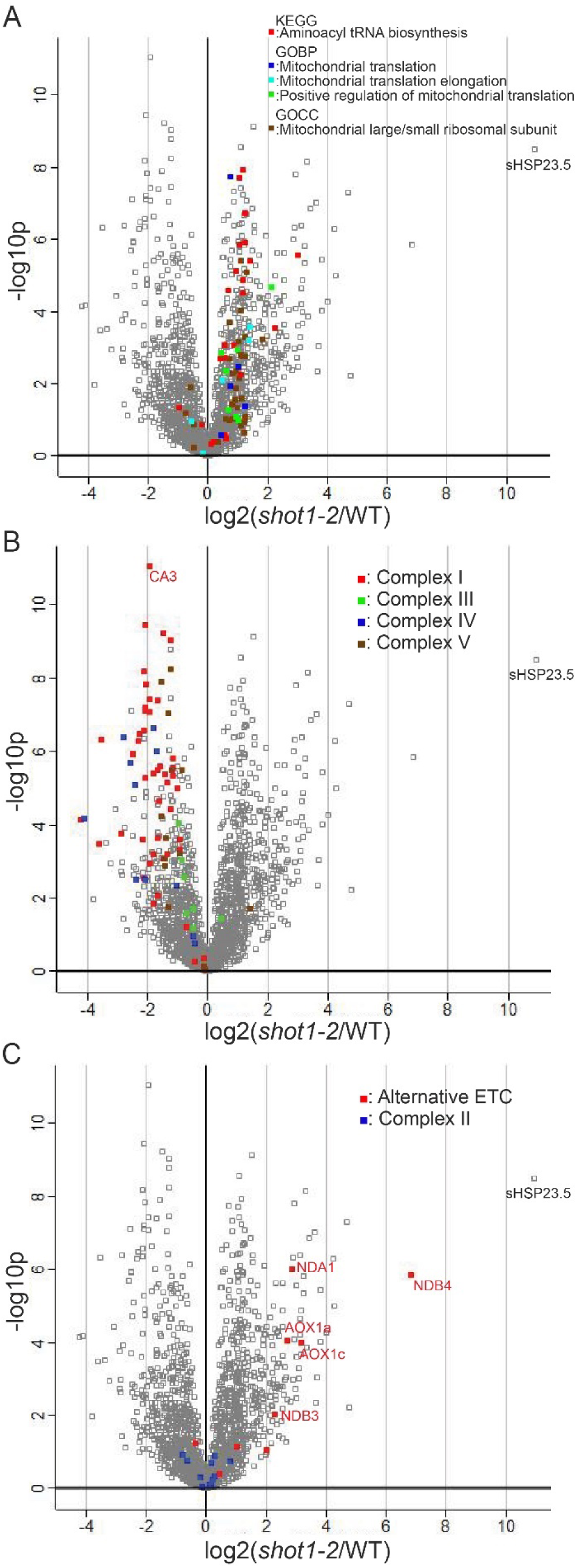
*shot1* has increased steady state levels of protein synthesis components, decreased complex I, III, IV and V subunits, but increased alternative electron transport components. Shown are volcano plots of 1988 proteins identified by mitochondrial proteomics, with colored squares for proteins in categories indicated (see Dataset S1). (A) Proteins related to translation are increased. (B) OXPHOS complexes containing mitochondria-encoded proteins are decreased. Both nucleus- and mitochondria-encoded proteins are labeled. (C) Complex II subunits, which are all nucleus-encoded are unchanged. Alternative ETC components are upregulated in *shot1*.

Proteins related to gene expression (GOBP: RNA modification; mRNA processing; Regulation of transcription, DNA-templated) were increased in the mutant (*SI Appendix*, Fig. S5B), consistent with the increase in mitochondria-encoded transcripts observed previously (23). Increased transcription and translation would also require proteins involved in mitochondrial gene expression such as PPR proteins, which were upregulated in the mutant (*SI Appendix*, Fig. S5C). Consistent with the slow growth of *shot1-2*, proteins involved in carbon metabolism (KEGG: Glycolysis/Gluconeogenesis; Pyruvate metabolism; Biosynthesis of amino acids) were mostly reduced (*SI Appendix*, Fig. S5D). In contrast, most tricarboxylic acid cycle enzymes did not change much in the mutant (*SI Appendix*, Fig. S5E).

In summary, proteomic analysis supports that the major defect in *shot1* mutants lies in the assembly of OXPHOS subunits encoded on the mitochondrial genome.

### SHOT1 interacts with mitochondrial DNA

To identify possible nucleic acid binding targets of SHOT1, C-terminal GFP fusion constructs under either a constitutive 35S promoter (*35S::SHOT1-GFP*) or the *SHOT1* native promoter (*SHOT1p::SHOT1-GFP*) were made. The transgenes were introduced into the *shot1-2* mutant background and verified to be functional based on recovery of *shot1* from growth defects (*SI Appendix*, Fig. S6). As a control, a transgenic line harboring mitochondria-targeted GFP (*Mito-GFP*) under the 35S promoter was used (29).

Multiple attempts to find RNA targets using crosslinking followed by immunoprecipitation were unsuccessful. We turned to finding DNA targets by immunoprecipitation of GFP crosslinked to DNA from isolated mitochondria of *Mito-GFP, 35S::SHOT1-GFP* and *SHOT1p::SHOT1-GFP* plants. Illumina sequencing of immunoprecipitated DNA revealed four enriched sites in *SHOT1-GFP* plants (*SI Appendix*, Table S1 and Fig. 4). All enriched peaks in *SHOT1-GFP* samples appear dispersed on the mitochondrial genome (GenBank: BK010421.1) (Fig. 4B) but are located upstream of different tRNA genes: *trnS*, t*rnM, trnG* and *trnF*.

**Fig. 4.**
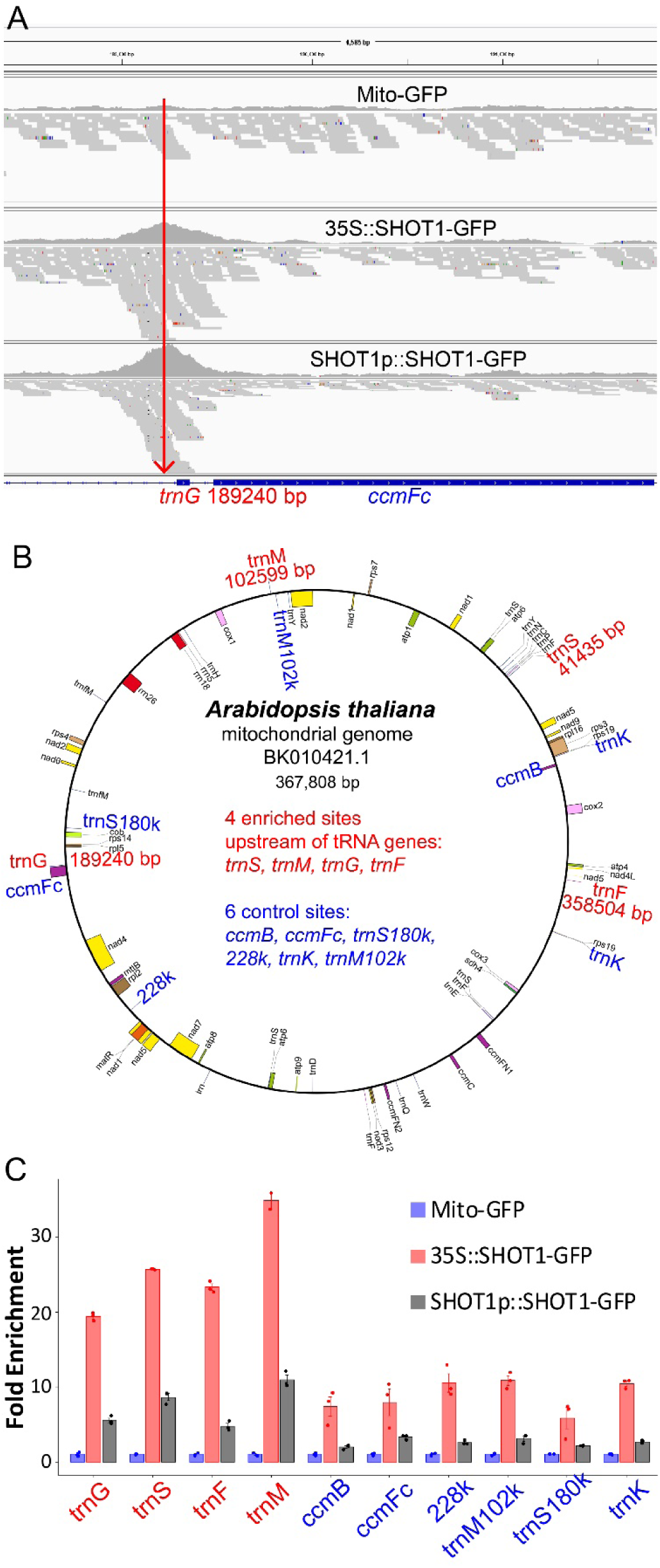
SHOT1-GFP binds to four sites on the mitochondrial genome. (A) An Integrative Genomics Viewer (IGV) snapshot of the site upstream of tRNA-Gly. (B) Location of the four enriched sites upstream of tRNA genes (red) and six control sites (blue). (C) Fold-enrichment of sites determined by qPCR following immunoprecipitation of DNA. Two biological replicates yielded similar results. The graph shows one data set; error bars = standard deviation of three technical replicates. Enrichment values were normalized to the average value of Mito-GFP samples, which was set to 1.0.

To validate the DNA-IP results, we designed qPCR primers surrounding the four tRNA peaks along with primers for six negative control sites. Control sites were *ccmB* and *ccmFC* genes (near *trnG*), one of the other five tRNA-Ser genes (*trnS180k*) near 180000 bp, tRNA-Lys genes (*trnK*) that are duplicates near *trnF*, ∼600 bp upstream region (*trnM102k*) of *trnM*, and a random site (228k) near 228,000 bp. Results were confirmed in DNA-IP-qPCR experiments on two independent biological replicates, with results from one replicate shown in Fig. 4C. As expected, the *SHOT1p::SHOT1-GFP* sample showed enrichment of the SHOT1 binding sites. *35S::SHOT1-GFP* samples had higher enrichment of the targets and slight enrichments of control sites that may indicate SHOT1 has weak, non-specific DNA binding activity that is detected when the protein is overexpressed.

Possible sequence-specificity of SHOT1 binding was examined using Multiple Em for Motif Elicitation (MEME) (30) with 102 bp sequences surrounding the enriched peaks. Motifs in the four SHOT1 binding sites are shown in *SI Appendix*, Fig. S7. A search of the first two motifs against the mitochondrial genome using FIMO (Find Individual Motif Occurrences) (31) returned only the four SHOT1 binding sites, suggesting SHOT1 binds these sites in a sequence specific manner.

Since SHOT1 binding sites are located immediately upstream of tRNA genes, we checked the possibility that SHOT1 regulated transcript levels of the four tRNA genes. RNA gel blot analysis was performed with two biological replicates of WT, *shot1-2*, and a complemented line (*SI Appendix*, Fig. S8). *trnS* and *trnF* transcript levels were higher, while those of *trnG* were lower in *shot1-2*, and *trnM* transcript levels were similar in all samples. The control transcripts, *trnK* and *trnQ*, were increased in *shot1-2*, which suggests that the overall increase in mitochondrial transcripts (23) also occurs at the level of tRNAs. The *ccmFC* transcript was reported to be lower in *shot1-2* (23), and *trnG*, which is co-transcribed with *ccmFC* (26) was similarly down-regulated. However, based on the proteomic data, *shot1-2* makes ccmFC protein at a slightly higher level than WT (Table 1), implying that *ccmFC* expression is post-transcriptionally regulated. Together, these results indicate that SHOT1 is not likely to affect OXPHOS biogenesis through regulation of these tRNA genes.

### SHOT1 interacts with AAA domain-containing proteins homologous to human ATAD3 proteins

Although SHOT1 clearly associates with mtDNA, our experiments do not point to a specific function in mitochondrial gene expression. Therefore, we investigated its possible function by identifying interacting proteins, using immunoprecipitation coupled with mass spectrometry (IP-MS). Three IP-MS experiments were performed with independent mitochondrial preparations from *35S::SHOT1-GFP* and *Mito-GFP* (a negative control) plants. As expected, immunoprecipitation with an anti-GFP nanobody (GFP-Trap, ChromoTek) consistently showed enrichment of SHOT1 in *35S::SHOT1-GFP* samples (Fig. 5A). Notably, three AAA family proteins were also enriched in all *35S::SHOT1-GFP* samples. To rule out artifacts from overexpression of SHOT1-GFP, a fourth experiment used GFP-Trap in an IP with mitochondria from *SHOT1p::SHOT1-GFP* plants in parallel to an IP from *35S::SHOT1-GFP* mitochondria using either GFP-Trap or blocked agarose beads (BAB, a negative control). The same AAA family proteins were enriched in both SHOT1-GFP samples with GFP-Trap, but not in the negative control (Fig. 5A). No other proteins were consistently immunoprecipitated in all *SHOT1-GFP* samples. The AAA family proteins identified are homologs of human <u>AT</u>Pase Family <u>A</u>AA <u>D</u>omain-Containing Protein 3 (ATAD3) proteins (32, 33). These *A. thaliana* ATAD3 proteins will be referred to as <u>S</u>HOT1 <u>B</u>inding <u>A</u>TPases (SBA) proteins. To confirm interaction between SHOT1-GFP and SBA1, we repeated the IP with mitochondrial lysates from transgenic plants carrying *SHOT1-GFP* controlled by either the 35S or native promoter. SBA1 was enriched in both samples (Fig. 5B). Importantly, recovery of SBA1 from plants carrying *SHOT1-GFP* driven by the native promoter rules out artifacts arising from over-expression of SHOT1-GFP.

**Fig. 5.**
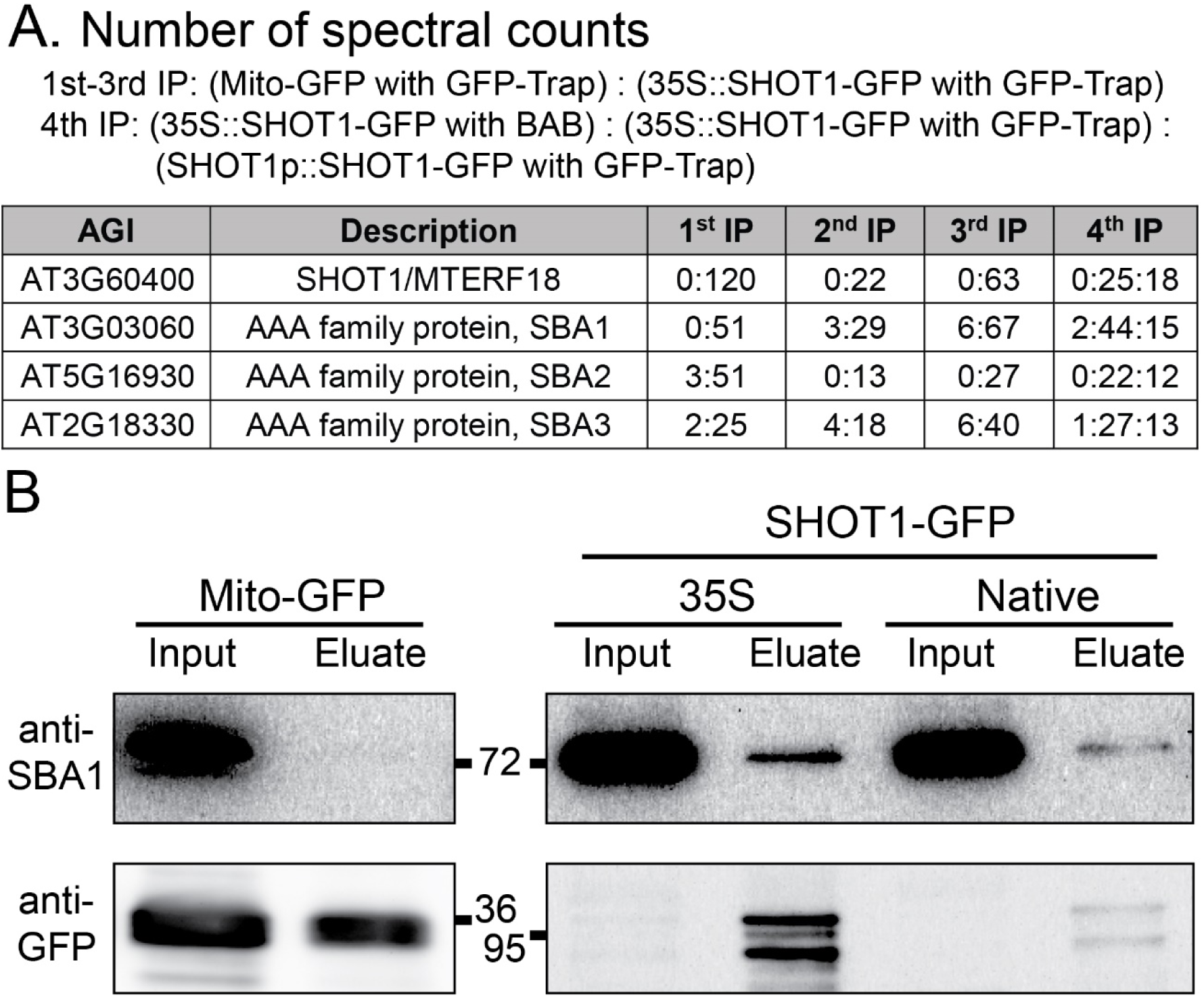
SHOT1-GFP interacts with SBA proteins. (A) Spectral counts showing enrichment of SBA proteins in SHOT1-GFP samples from experiments using four independent mitochondrial preparations. Three IPs were performed using GFP-Trap agarose beads with mitochondria from Mito-GFP or 35S::SHOT1-GFP. A fourth IP was performed with 35S::SHOT1-GFP mitochondria using either blocked agarose beads (BAB, the control) or GFP-Trap beads, and with SHOT1pro::SHOT1-GFP mitochondria using GFP-Trap beads. (B) Immunoblot analysis shows SBA1 is enriched in SHOT1-GFP immunoprecipitates. Mitochondrial lysates from Mito-GFP, SHOT1-GFP under either 35S or a native promoter were immunoprecipitated with GFP-Trap. 1 % of total input and 50% of total eluate were separated for immunoblot analysis. The immunoblot was first probed with anti-SBA1 antibody and then with anti-GFP antibody. Strong Mito-GFP (anti-GFP) signals have been lowered using image analysis software.

Most animals have one ATAD3 gene with the exception of primates, which have two additional paralogs, ATAD3B and ATAD3C, likely derived from recent duplication events (34). *A. thaliana* has four ATAD3 proteins all on different chromosomes, three of which (SBA1, 2 and 3) were detected in our experiments. The four SBA proteins represent two clades: SBA1 (AT3G03060) and SBA2 (AT5G16930) with ∼86 % protein identity, and SBA3 (AT2G18330) and SBA4 (AT4G36580) with ∼85% protein identity (*SI Appendix*, Fig. S9A). ATAD3 proteins including these SBAs have two predicted central transmembrane (TM) domains, which could span the mitochondrial inner and outer membrane (35). The C-terminus-containing AAA domain resides in the matrix, while the N-terminus is reported to face outside the mitochondrion (36). All four SBA proteins were detected in our mitochondrial proteomics data (Dataset S1) and in recently published data (37), but not in publicly available chloroplast proteomics studies, consistent with exclusive mitochondrial localization. ATAD3 proteins are conserved between plants and animals (35% identity and 52% similarity between human ATAD3A and SBA1) (*SI Appendix*, Fig. S9B), indicating they play fundamental roles in mitochondria.

### SHOT1 is required for mitochondrial nucleoid organization

Animal ATAD3 proteins appear to play critical roles in organization of mitochondrial nucleoids (32, 33, 38–40). Our findings that SHOT1 interacts with both mitochondrial DNA and ATAD3/SBA proteins suggested that SHOT1 might also be involved in mitochondrial nucleoid organization. In addition, increased DNA copy number (23) and higher accumulation of nucleoid proteins (*SI Appendix*, Fig. S5F) in *shot1-2* may be a compensatory response resulting from defective nucleoids. Therefore, we compared mitochondrial nucleoids between WT and *shot1* plants after staining nucleoids with PicoGreen. The *ndufs4* mutant (a T-DNA insertion mutant in AT5G67590), which has a mutation in a complex I subunit (41), was included as a control that was not expected to have nucleoid defects. Nucleoids of WT and *ndufs4* mitochondria formed tight, intense foci inside mitochondria (Fig. 6A). However, nucleoid signals in *shot1-1* and *shot1-2* mitochondria were weaker generally, enlarged and dispersed (Fig. 6A and B), similar to the nucleoid phenotype of human cells with reduced ATAD3A (40). In addition to nucleoid abnormalities, *shot1* mitochondria appeared more spherical and larger than WT, and we frequently observed giant mitochondria (diameter up to 3 µm) with a relatively stronger nucleoid signal in *shot1* mutants. Nucleoids of *ndufs4* were indistinguishable from those of WT mitochondria implying that disturbed mitochondrial nucleoids in *shot1* are not likely caused solely by defects in complex I.

**Fig. 6.**
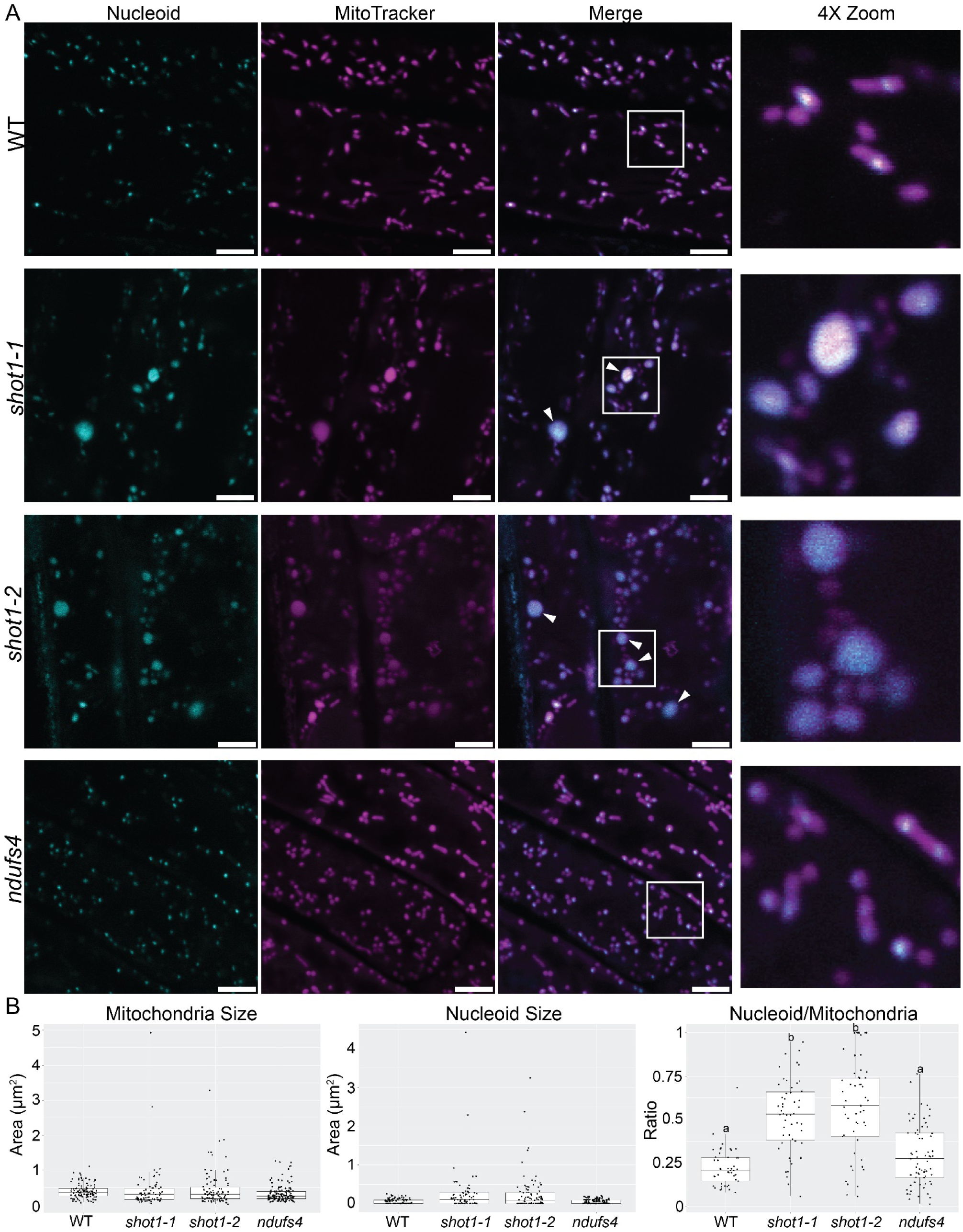
*shot1* mutants have diffuse and enlarged mitochondrial nucleoids. (A) Mitochondria from root epidermal cells stained with PicoGreen (for nucleoids) and MitoTracker Orange. Arrowheads indicate giant mitochondria. Boxed areas are shown at higher magnification. Scale bars = 5 µm. (B) Images in (A) were processed using General Analysis 3 in the NIS-Elements package (Nikon). Nucleoid sizes relative to mitochondrial sizes are plotted in the third panel, after removal of mitochondria without nucleoids. Different letters indicate significant differences (p<0.01) by one-way ANOVA followed by Tukey’s post hoc test.

The possible role of SHOT1 in mitochondrial nucleoid organization led us to examine whether SHOT1-GFP localizes as expected for a nucleoid protein. In plants, mtDNA can be visualized as punctate structures (42) and is thought to be packaged into nucleoids similar to mammalian mtDNA (43). While Mito-GFP appears as diffuse GFP signals overlapping with MitoTracker Orange (Fig. 7A), distinctive compact GFP signals inside mitochondria, reminiscent of mitochondrial nucleoids, were observed for SHOT1-GFP (Fig.7B and 7C). Thus, our data suggest that SHOT1 function is required for proper organization of mitochondrial nucleoids, as well as normal mitochondrial morphology.

**Fig. 7.**
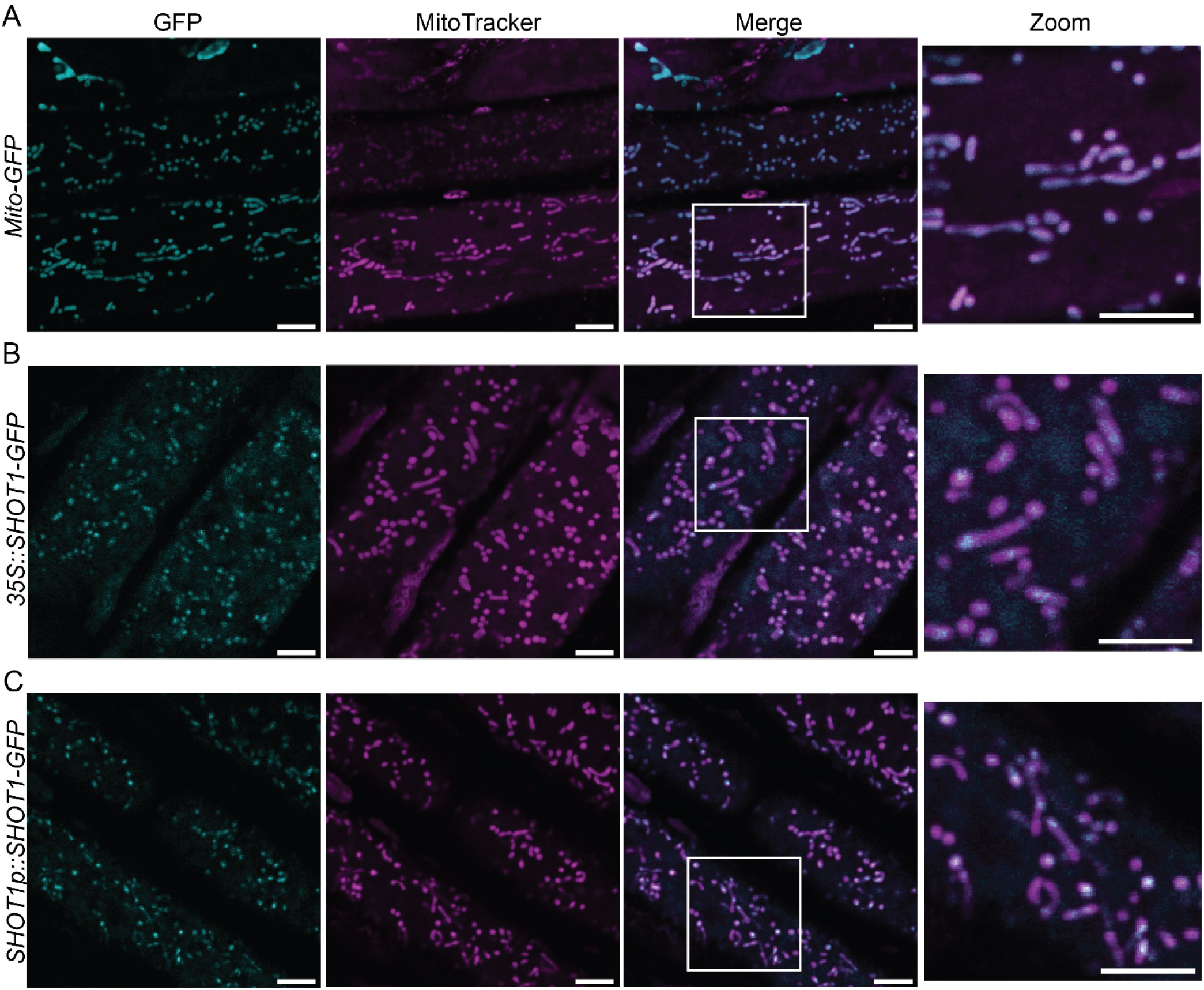
SHOT1-GFP localizes as expected for mitochondrial nucleoids. (A) *Mito-GFP*, (B) *35S::SHOT1-GFP* and (C) *SHOT1p::SHOT1-GFP* seedlings. Boxed area shown at higher magnification. Scale bars represent 5 µm.

### Disruption of SBA1 function leads to disturbed nucleoid organization and enhanced thermotolerance

The fact that SHOT1 appears required for proper mitochondrial nucleoid organization and interacts with SBA proteins, whose animal ATAD3 homologs are also implicated in nucleoid organization, led us to hypothesize that SBA mutants should show altered nucleoid organization and could exhibit other *shot1* phenotypes. T-DNA insertion mutants that disrupt gene expression of each *SBA* gene were obtained (*sba1-1*: GK-217D03, *sba1-2*: GK-488F07, *sba2-1*: SAIL_1215_E01, *sba2-2*: SALK_140274, *sba3-1*: SALK_007874, *sba4-1*: SALKseq_127403.1, *sba4-2*: SALK_006588) (*SI Appendix*, Fig. S10A and B). All single homozygous SBA mutants grew like WT, which was not surprising given their sequence similarity within the two SBA clades (SBA1 and 2, vs. SBA3 and 4). However, we were unable to recover a double mutant of *sba1-1* with *sba3-1*, indicating these SBA proteins are essential for plant growth as are ATAD3 proteins in animals (44, 45).

Localization of the SBA1 protein was examined with a C-terminal GFP fusion driven by the native SBA1 promoter and transformed into *sba1-1 sba3-1/+* mutant plants. In the T3 generation, we identified SBA1-GFP plants in both the *sba1-1* single mutant and *sba1-1 sba3-1* double mutant backgrounds. Interestingly, both genotypes showed retarded growth, with the double mutant having a more severe phenotype (*SI Appendix*, Fig. S10C). Recovery of SBA1-GFP plants in *sba1-1 sba3-1* double mutant background indicates that the SBA1-GFP fusion protein is partially functional.

However, the growth defects, even in the *sba1-1* single mutant carrying the SBA1-GFP transgene indicate that the C-terminal GFP tag interferes with the function of not only SBA1, but also of SBA3, possibly acting in a dominant-negative fashion. The human ATAD3A protein has been shown to form hexamers typical of AAA+ proteins (35). Lethality of the double *sba1 sba3* mutant and phenotype of SBA1-GFP transgenic plants suggest SBA proteins form hetero-hexamers.

We examined mitochondria of SBA1-GFP plants to localize the protein and visualize mitochondrial nucleoids. GFP signal was detected mainly at the perimeter of mitochondria, confirming SBA1 membrane localization. Notably, most mitochondria appeared more spherical than in WT and giant mitochondria were observed, as seen in *shot1* mutants (Fig. 8A). We then visualized mitochondrial nucleoids with PicoGreen. Although the PicoGreen signal cannot be distinguished spectrally from SBA1-GFP, the signals are separated in matrix versus membrane, respectively. When compared with GFP signals without PicoGreen staining (Fig. 8A), PicoGreen/GFP signals in SBA1-GFP mitochondria were diffuse inside the mitochondrial matrix in both *sba1-1* and *sba1-1 sba3-1* backgrounds, indicating that nucleoid organization is disrupted when SBA function is compromised (Fig. 8B and *SI Appendix*, Fig. S10B).

**Fig. 8.**
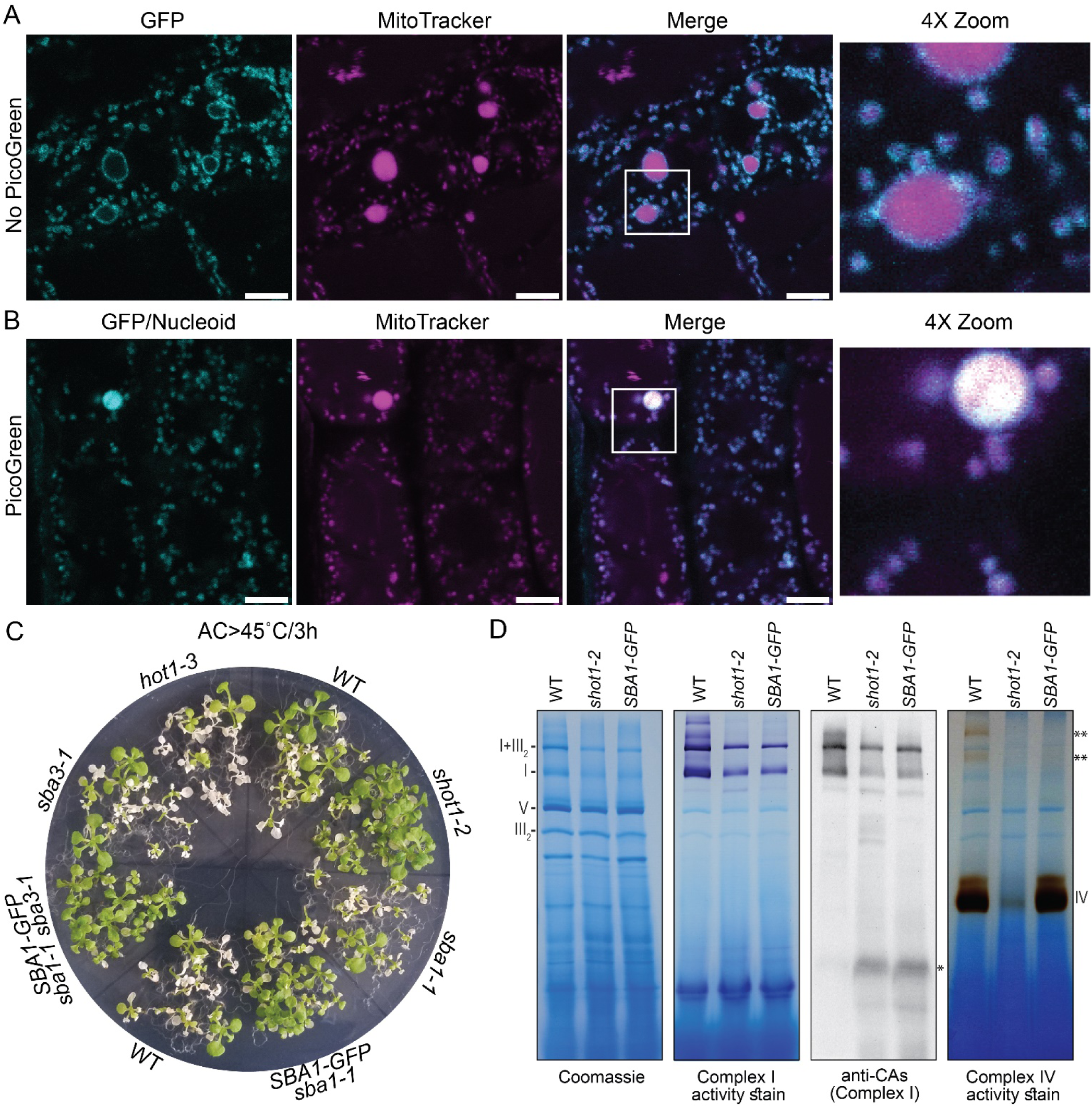
*SBA1-GFP* plants have disturbed mitochondrial nucleoid organization and are more tolerant to heat stress. Root epidermal mitochondria of *SBA1-GFP* plants (*sba1-1 sba3-1* background) were observed without (A) or with (B) PicoGreen staining. Boxed areas shown at higher magnification. Scale bars = 5 µm. (C) 11-day-old seedlings were heat-stressed for 3 h at 45 °C after acclimation (38 °C/1.5h plus 2h at room temperature) and photographed 8 days later. (D) BN-PAGE of mitochondria from wild type, *shot1-2* and *SBA1-GFP* (*sba1-1 sba3-1* background) plants. Gels were Coomassie-stained or stained for NADH oxidase (complex I) or cytochrome oxidase (complex IV) activity. Immunoblot with anti-CAs detect complex I and assembly intermediates. Complexes I, III_2_, IV and V and supercomplex I+III_2_ are labelled. Asterisk indicates an early assembly intermediate of complex I. Double asterisks indicate supercomplexes containing complex IV.

The stark similarities in mitochondrial morphology and nucleoid organization between *shot1* mutants and *SBA1-GFP* plants combined with the physical interaction of SHOT1 and SBA1 strongly suggest that they function together in mitochondria. Therefore, *SBA1-GFP* plants were expected to show higher thermotolerance as do *shot1* mutants. We subjected *SBA1-GFP* plants in parallel with *sba1-1, sba3-1* and *hot1-3* (HSP101 null mutant) plants to 45 °C heat stress for 3 hours after acclimation treatment (38 °C for 1.5h followed by 2h at 22 °C). As predicted, plants carrying *SBA1-GFP* in either the *sba1-1* or *sba1-1 sba3-1* backgrounds were more heat-tolerant than WT, similar to *shot1-2* plants (Fig. 8C). We then examined OXPHOS complexes in *SBA1-GFP* (*sba1-1 sba3-1* background) plants by BN-PAGE, followed by complex I and IV activity staining and immunoblot analysis of complex I. Similar to *shot1-2, SBA1-GFP* plants exhibited a complex I defect (Fig. 8D). However, complexes III and V appear similar to WT, and complex IV activity shows little difference between WT and *SBA1-GFP*, although activity is low in super complexes (double asterisks in Fig. 8D), likely due to reduced complex I. The capacity of *SBA1-GFP*, but not *shot1-2*, to assemble complex IV might derive from the only partial disruption of SBA1 function by the GFP tag. Alternatively, SHOT1 might have SBA1-dependent and -independent functions, with nucleoid organization and complex I accumulation requiring, and complex IV accumulation not requiring interaction with SBA1.

In summary, our data indicate that SHOT1 in cooperation with the conserved ATAD3/SBA proteins, play essential roles in mitochondrial nucleoid organization and OXPHOS assembly in plants.

## Discussion

Work presented here identifies the mTERF family protein SHOT1 as binding mitochondrial DNA at four sites upstream of specific tRNA genes. However, we found no evidence that it regulates expression of these genes directly. Instead, SHOT1 appears involved in nucleoid organization, as evidenced by the loss of discrete nucleoid foci in *shot1* mitochondria and interaction of SHOT1 with SBA1, 2 and 3, which are ATAD3 homologs that are known players in mitochondrial nucleoid organization in animals (32, 33, 38). In mammalian mitochondria, mitochondrial transcription factor A (TFAM) is important for packaging mtDNA into nucleoids (46, 47). Mammalian nucleoid preparations also contain DNA replication and transcription components, RNA binding proteins and some ribosomal proteins, indicating RNA processing and ribosome assembly occur in proximity to nucleoids (32, 38, 48, 49). Plants lack a TFAM homologue, perhaps reflecting the divergence of mtDNA maintenance and mitochondrial gene expression control between plants and animals (4); other components of plant mitochondrial nucleoids are poorly defined (50, 51). Despite these known differences in nucleoids, our data document a conserved function for ATAD3 homologues in maintaining nucleoid structure across kingdoms.

We observed previously that *shot1-2* has increased mtDNA and transcripts (23). Here we document an increase in translation in mutant mitochondria, but dramatically reduced levels of mitochondria-encoded OXPHOS subunits. Nucleus-encoded complexes I, III, IV and V subunits were also significantly reduced in *shot1-2*. These data are in stark contrast to unchanged levels of complex II subunits, all of which are nucleus-encoded. The most plausible explanation is that the *shot1* mutants are defective in assembly of mitochondria-encoded OXPHOS subunits. Assembly failure would trigger degradation of unassembled nucleus-encoded OXPHOS subunits by the mitochondrial protein quality control system, components of which are increased in *shot1-2*. These defects are likely communicated with the nucleus, leading to an increase of alternative respiratory chain components, such as AOX and alternative NADH dehydrogenases, which would contribute to lowering ROS levels and enhancing thermotolerance in *shot1* mutants.

Our findings strongly indicate that the primary defect of *shot1* mutants is organization of mitochondrial nucleoids. Disrupted nucleoids in *shot1* mutants appear to affect proper assembly of mitochondria-encoded OXPHOS subunits, triggering many adjustments via retrograde signaling, including increasing mtDNA levels, transcript levels and translation. These changes are no doubt critical to the enhanced heat tolerance of *shot1*. The fact that plants with disrupted SBA1 function also show disrupted nucleoids and enhanced heat tolerance further indicates specific signaling of the nucleoid status to the rest of the cell; other mutants defective in OXPHOS are not heat tolerant (23). The mechanism by which OXPHOS assembly, in particular complex I function, is affected in *shot1* mutants and SBA1-defective plants is not clear. However, our data suggest a close connection between nucleoid organization and OXPHOS assembly. mtDNA is tightly associated with the mitochondrial inner membrane in eukaryotes (43, 52, 53), but in *shot1* mutants mtDNA appears distributed throughout the matrix rather than tethered to a specific inner membrane location. Translation and membrane insertion of the hydrophobic, mitochondria-encoded OXPHOS subunits may need to occur near the membrane, which could require a membrane-associated nucleoid. Although mitochondrial transcription and translation are not documented to be coupled, nucleoid organization and membrane tethering could possibly facilitate membrane insertion of the hydrophobic OXPHOS subunits.

Although several plastid-targeted mTERFs are seen associated with nucleoids (11, 54–56), mitochondria-targeted mTERFs have not been reported to be nucleoid-associated in plants. Human mTERF1 was identified among a number of ATAD3A-interacting proteins (38), although the significance of this interaction has not been studied. ATAD3 proteins are not likely to interact directly with mtDNA as they were not detected in nucleoid preparations after crosslinking (32). It is possible that mTERFs mediate the interaction between mtDNA and ATAD3 proteins. SHOT1 is, however, in very low abundance (undetectable in our mitochondrial proteomics experiments) compared to SBA proteins, suggesting that SHOT1 is not a structural component of mitochondrial nucleoids as is TFAM in human mitochondria.

While the matrix-located, C-terminal ATPase domain of animal ATAD3 proteins interacts with mitochondrial nucleoids and ribosomes, the N-terminal domain appears to interact with cytosolic proteins (38, 57). A cytosolic dynamin-related GTPase protein, DRP1, which oligomerizes as part of the mitochondrial fission machinery, was recently reported to interact with ATAD3A in humans (39). Mitochondrial fission events occur in close proximity to mtDNA replication sites and to ER-mitochondria contact sites in yeast and mammalian cells (58). From these observations, we speculate that ATAD3 proteins communicate the status of mitochondrial nucleoids to the ER and to the DRP1 fission machinery (DRP3a (AT4G33650) and DRP3b (AT2G14120) in *A. thaliana*). When mitochondrial nucleoids are not properly organized, as in *shot1* mutants or *SBA1-GFP* plants, mitochondrial fission may be compromised. This would result in the observed giant mitochondria, similar to the giant mitochondria, along with elongated mitochondria, that are observed in *drp3a drp3b* mutants (59).

In addition to their role in mitochondrial nucleoid organization, ATAD3 proteins in animals are associated with diverse cellular functions including mitochondrial dynamics, protein synthesis, lipid and steroid biosynthesis, and mitophagy (36, 38, 60–62). How this protein is involved in these many different processes is unknown, but all are associated with functions of protein complexes at the mitochondria-ER interface (35). Considering the conservation of ATAD3 proteins in multicellular eukaryotes, SBA proteins are likely to perform similar functions in plants. Although the mTERF protein family is also unique to multicellular eukaryotes, it has dramatically diverged and expanded in plants, potentially reflecting the evolution of complex genomes and gene expression systems in plant mitochondria. Our study emphasizes the importance of understanding the role of mitochondrial nucleoid organization in the assembly of OXPHOS machinery, which is essential for supplying energy to plants and food for humanity. How nucleoid organization may then be linked to the function of mitochondria-ER contact sites, retrograde signaling and enhanced plant heat tolerance are novel areas for further research.

## Materials and Methods

The materials and methods are described in *SI Appendix, SI Materials and Methods*.

## Acknowledgements

The authors acknowledge services of the University of Massachusetts Amherst Institute of Applied Life Sciences Cores: the Genomics Resource Laboratory, Mass Spectrometry Center and Light Microscopy Facility, as well as the University of Massachusetts Medical School Mass Spectrometry Facility for mitochondrial proteomics, and Kirk Mackinnon for help analyzing DNA-IP-Seq. M.K. and E.V. thank Tianxiang Liu for help with mitochondrial preps and Patrick Treffon for GFP-Trap beads and advice. Supported by National Science Foundation grant IOS 1354960 to E. V. and Deutsche Forschungsgemeinschaft grant 400681449/GRK2498 to K.K..

## Footnote

Minsoo Kim: Designed and performed research, analyzed data and wrote the paper; Vincent Schulz, Lea Brings and Theresa Schoeller: performed research; Kristina Kühn and Elizabeth Vierling: designed research, analyzed data, wrote the paper.

