## Supplementary material for "Mitochondrial nucleoid organization and biogenesis of complex I require mTERF18/SHOT1 and ATAD3 in *Arabidopsis thaliana*": SI Appendix

**ORCID**:

M.K. 0000-0002-9997-6489

K.K. 0000-0001-6747-783X

E.V. 0000-0002-0066-4881

**Corresponding Author:** Elizabeth Vierling

**This PDF file includes:**

Supplementary text

Figures S1 to S10

Tables S1 to S2

Legends for Datasets S1

SI References

**Other supplementary materials for this manuscript include the following:**

Datasets S1

Supplementary Information Text

Supplemental Material and Methods:

**Plant Material and Growth Conditions.** *Arabidopsis thaliana* mutants, *shot1-1* (G105D), *shot1-2* (GK_040C09) and *ndufs4* (SAIL_596_E11) are in the wild type Col-0 background. *SHOT1* genomic DNA complemented *shot1-2* line has been described previously (1). Seeds were surface sterilized and plated on 0.5× Murashige and Skoog (MS) medium (2) supplemented with 0.5% sucrose and 0.8% phytoagar. Plants were grown in controlled growth chambers at 22°C/18°C, 16-h-day/8-h-night cycle with fluorescent lamps (100 μmol m^–2^ s^–1^). Seedlings for mitochondria isolation were grown in 0.5x MS liquid media containing 0.5% sucrose with shaking in the same growth conditions. Mitochondria-targeted GFP (Mito-GFP) seeds were obtained from Dr. David Logan (3).

**Generation of Constructs and Transgenic Lines.** To make the SHOT1-GFP construct under the 35S promoter, the coding sequence of *SHOT1* was amplified with the primers SHOT1-1, 5’-CACCATGTTCATGGTTCGATTGAAATT-3’ and SHOT1-R1, 5’-GGATACAGTGTCTCTGGTTTTTCT-3’ and cloned into pENTR/D-TOPO (Invitrogen). The coding sequence was then transferred to the pMDC83 binary vector under 2X 35S promoter (4) by Gateway LR cloning (Invitrogen).

To generate the SHOT1-GFP construct under the native promoter, the 4649 bp SHOT1 genomic DNA fragment containing the promoter (2971 bp region upstream of the start codon) and the ORF without the stop codon was amplified with the primers SHOT1-F1, 5’-CACCACAAGAAGAGGTGGAGTGGG-3’ and SHOT1-R1 and cloned into pENTR/D-TOPO (Invitrogen). The sequence was then transferred to the pMDC107 binary vector (4) by Gateway LR cloning (Invitrogen). The constructs were introduced into *shot1-2* plants by agrobacterium mediated transformation with the floral dip method (5).

For translational fusion of GFP to the C-terminus of SBA1, the 5058 bp SBA1 genomic DNA fragment containing the promoter (1824bp region upstream of the start codon) and the ORF without the stop codon was amplified with the primers SBA1-1, 5’-CACCCATCCCGGATTCTGCAGTTG-3’ and SBA1-2, 5’-TTTTGAATCGACTCCAGCCAAT-3’ and cloned into pCR8/GW/TOPO (Invitrogen). The fragment was then transferred to the pMDC107 binary vector by Gateway LR cloning (Invitrogen). The construct was introduced into *sba1-1 sba3-1/+* plants by agrobacterium mediated transformation with the floral dip method (5).

**Isolation of Mitochondria.** Surface sterilized seeds were grown in 0.5x MS liquid media containing 0.5% sucrose until stages 1.02-1.04 (6) or on soil until stage 6.00 (6) in a growth chamber at 22°C/18°C, 16-h-day/8-h-night cycle with fluorescent lamps (100 μmol m^–2^ s^–1^). 15-30 grams of whole seedlings grown in liquid media or 20-60 grams of aerial parts of plants grown on soil were harvested and ground at 4°C in ~30 ml extraction buffer (0.3 M sucrose, 25 mM K_4_P_2_O_7_, 10 mM KH_2_PO_4_, 2 mM EDTA, 1% [w/v] PVP-40, 1% [w/v] BSA, 5 mM cysteine, and 25 mM sodium ascorbate, pH7.5) per 10 grams of plant material using a mortar and a pestle. The homogenate was spun at 1700g for 5 min and the supernatant was spun at 20000g for 10 min. The pellet was resuspended in wash buffer (0.3 M sucrose, 1 mM EGTA, and 10 mM MOPS/KOH, pH 7.2) and subjected to additional low (2500g)- and high (20000g)-speed centrifugations. The pellet was resuspended in a small volume (around 1 ml) of wash buffer and loaded on top of a 0-4.4% PVP-40 gradient in 28% Percoll, 0.3 M sucrose, 10 mM MOPS/KOH, pH7.2. The gradient was centrifuged at 40000g for 45 min. The mitochondrial fraction on the bottom of the gradient was collected and washed three times by centrifugation at 30000g for 10 min and resuspension of the pellet in wash buffer. The purified mitochondrial fraction was saved as a mitochondrial suspension at -80°C.

**Blue Native PAGE (BN-PAGE), immunodetection of protein complexes and in-gel activity staining**

Mitochondrial proteins (100 µg for Coomassie staining and in-gel activity staining; 50 µg for immunolabeling of complexes) were solubilized in digitonin solution (5% digitonin, 30 mM HEPES, 150 mM potassium acetate, 10% glycerol) and prepared for electrophoresis as described (7). Protein complexes were separated in a 4.5 to 16% gradient gel (8). Following migration, protein complexes were transferred onto a polyvinylidene difluoride (PVDF) membrane (GE Healthcare) in cathode buffer (50 mM Tricine, 15 mM Bis-Tris-HCl, pH 7.0) for 16 h at 4°C. Alternatively, gels were Coomassie-stained or subjected to in-gel activity staining (see below). Membranes were Coomassie-stained to control for efficient protein transfer and equal loading, destained and probed with antibodies against CA2 (9), the mitochondrial Rieske iron-sulfur protein (RISP) (10), Cox2 (AS04 053A; Agrisera) or ATP2 (AS05 085; Agrisera). Signal detection was performed using Clarity Western ECL Blotting Substrate (Bio-Rad).

In-gel activity staining for NADH oxidase activity (complex I) or cytochrome oxidase activity (complex IV) was carried out as described (11).

**SDS-PAGE and Immunoblot Analysis.** SDS-PAGE and immunoblot analysis were performed as described previously (1). Rabbit anti-SBA1 antibody was generated against the recombinant protein purified from *E. coli* expressing the C-terminal mitochondrial matrix domain of the protein (300aa-628aa) by Pocono Rabbit Farm and Laboratory. Antibody dilutions were 1:4000 for antibodies against SBA1 and GFP (Roche #11814460001); 1:5000 for antibodies against Nad1 (12), Nad6 (13), Cob (12), RISP (10), Cyt c1 (Gift from Gottfried Schatz; Basel University), Cox1 (12), Cox2 (AS04 053A; Agrisera), ATP2 (AS05 085; Agrisera), PDC E1alpha (Gift from Thomas E. Elthon; University of Nebraska), MnSOD (AS09 524; Agrisera) and AOX (14); 1:10000 for the antibody against PORIN (Gift from Thomas E. Elthon; University of Nebraska); 1:20000 for the antibody against CA2 (9); 1:50000 for antibody against Nad9 (15); 1:200000 for anti-rabbit IgG horseradish peroxidase conjugated (GE Healthcare); 1:50000 for anti-mouse IgG (SantaCruz). Thermo Scientific SuperSignal West Femto Maximum Sensitivity ECL Substrate was used to visualize the signals with a G:Box iChemi XT(Syngene).

**RNA isolation and RNA gel-blot hybridization**

RNA was extracted from seedlings grown on MS-agar until stage 1.02 (6) using TRIsure (Bioline). Total RNA (3 µg) was resolved on 1.2% w/v agarose/formaldehyde gels and transferred onto positively charged nylon membranes (Roche Life Science), followed by membrane staining with methylene blue (0.04% (w/v) in 0.5M sodium acetate, pH 5.2). Digoxigenin-labelled DNA probes were generated using Arabidopsis cDNA as template, the PCR DIG probe synthesis kit (Roche Life Science), and primer pairs listed in *SI Appendix*, Table S2. Hybridizations were performed in DIG Easy Hyb solution (Roche Life Science) according to the manufacturer’s instructions; chemiluminescent detection of signals was performed using anti-digoxigenin-alkaline phosphatase conjugates and CSPD reagent (Roche Life Science).

**Analysis of transcript editing**

Total RNA was depleted of contaminating genomic DNA by treatment with RQ1 RNase-free DNase (Promega) and confirmed by PCR to be free of detectable amounts of DNA. To produce cDNA, 1 µg of DNA-free RNA was reverse-transcribed with RevertAid H Minus Reverse Transcriptase (ThermoFisher Scientific) according to the manufacturer’s protocol using the gene-specific primers listed in *SI Appendix*, Table S2. cDNA fragments were PCR-amplified and sent for Sanger sequencing at Microsynth Seqlab GmbH.

**Polysome analysis.** Polysome fractionation was done according to (16); one third of every fraction was used for RNA gel blot hybridization.

**Mitochondrial Proteomics.** Wild-type Columbia-0 and *shot1-2* were grown in 0.5X MS liquid media supplemented with 0.5% sucrose until stages 1.02-1.04 (6). 40 micrograms of mitochondrial proteins (three biological replicates with three technical replicates each) were run on an SDS-PAGE gel. Downstream analyses were performed at the mass spectrometry facility at the University of Massachusetts Medical School. The entire protein region of the gel was excised and subjected to in-gel trypsin (Promega V511A) digestion after reduction with DTT and alkylation with iodoacetamide. Peptides eluted from the gel were lyophilized and re-suspended in 25µL of 5% acetonitrile (0.1% (v/v) TFA). A 3µL injection was separated on a Waters NanoAcquity UPLC in 5% acetonitrile (0.1% formic acid) at 4.0 µL/min for 4.0 min onto a 100 µm I.D. fused-silica pre-column packed with 2 cm of 5 µm (200Å) Magic C18AQ (Bruker-Michrom). Peptides were eluted at 300 nL/min from a 75 µm I.D. gravity-pulled analytical column packed with 25 cm of 3 µm (100Å) Magic C18AQ particles using a linear gradient from 5-35% of mobile phase B (acetonitrile + 0.1% formic acid) in mobile phase A (water + 0.1% formic acid). Ions were introduced by positive electrospray ionization via liquid junction at 1.5kV into a Thermo Scientific Q Exactive hybrid mass spectrometer. Mass spectra were acquired over m/z 300-1750 at 70,000 resolution (m/z 200) with an AGC (Automatic Gain Control) target of 1e6, and data-dependent acquisition selected the top 10 most abundant precursor ions for tandem mass spectrometry by HCD fragmentation using an isolation width of 1.6 Da. Peptides were fragmented by a normalized collisional energy of 27, and fragment spectra acquired at a resolution of 17,500 (m/z 200).

Raw data files were processed with MaxQuant (version 1.6.8.0) against the *Arabidopsis thaliana* (Uniprot) FASTA file (downloaded 06/2019). Search parameters included Trypsin/P specificity, up to 2 missed cleavages, a fixed modification of carbamidomethyl cysteine, and variable modifications of oxidized methionine, and N-terminal acetylation. Label free quantification was done using the MaxLFQ method (17). Downstream statistical analysis was performed using Perseus (version 1.6.10.0). Protein groups detected in at least five runs in either *shot1-2* or WT groups (9 runs in each group) were filtered resulting in 1998 total protein groups. Raw data were deposited to MassIVE repository (MSVxxxx).

**Immunoprecipitation of Mitochondrial DNA, Library Construction and Sequencing.** Mitochondria (500 µg mitochondrial proteins) from Mito-GFP, 35S::SHOT1-GFP, and SHOT1pro::SHOT1-GFP were isolated, resuspended in 1ml cross-linking buffer (0.5 M sucrose, 20 mM HEPES pH7.5, 2 mM EDTA, 7 mM β-mercaptoethanol, 1 % formaldehyde) and incubated for 30 min on ice. The cross-linking was quenched with 133 mM glycine and spun down at 16000 g for 20 min. Mitochondrial pellets were resuspended with 1 ml IP buffer (50 mM HEPES pH 7.5, 150 mM NaCl, 5 mM MgCl_2_, 0.5% Nonidet-P40, 0.5% Sodium deoxycholate, 0.05% SDS, complete protease inhibitor cocktail (Sigma P9599)) by pipetting up and down on ice. Samples were sonicated for 6 cycles of 10 sec on/50 sec off at 40% power (Sonics and Materials #VC505) and spun down at 16000g for 10 min to remove debris. 10 µl was set aside as input DNA. The remaining supernatants were precleared by incubating for 1 hr at 4 °C with blocked magnetic agarose beads (ChromoTek #bmab-20) equilibrated with IP buffer. Precleared samples were then incubated for 1 hr at 4 °C with GFP-Trap (ChromoTek #gtma-20) equilibrated with IP buffer. The beads were washed five times with 1 ml IP buffer before adding 100 µl elution buffer (0.2 M glycine, 0.5 M NaCl, 0.05% Tween-20, pH2.5) and incubating for 1 min at 37 °C. After transferring eluates to a new tube, 50 µl of 1 M Tris-HCl pH9.3 was added to neutralize eluates. The elution step was repeated twice more resulting in 450 µl total eluate volume.

To reverse cross-link, Proteinase K was added at 0.5 mg/ml to the eluates and incubated overnight at 37 °C. 440 µl of TE was added to the input DNA samples and treated in the same manner with proteinase K and overnight incubation at 37 °C. The samples were combined with a second aliquot of proteinase K and incubated at 65 °C for at least 6 hr. DNA samples were ethanol-precipitated overnight at -20 °C, resuspended in 100 ul sterile water and stored at -20 °C until later use.

Immunoprecipitated DNA samples were used to make libraries for sequencing following the manufacturer’s instructions (ACCEL-NGS 2S Plus DNA library kits, Swift Biosciences #21024). 151bp paired-end sequencing was performed using Illumina MiSeq at the Genomics Resource Laboratory, University of Massachusetts Amherst.

Sequencing data are available under GEO accession record GSE150262.

**Sequencing Data Processing and Analysis.** The sequencing data were uploaded to the Galaxy web platform, and we used the public server at usegalaxy.org to analyze the data (18). Sequencing reads were trimmed with Trim Galore and mapped against the Arabidopsis mitochondrial genome (GenBank: BK010421) using Bowtie2 (version 2.3.2). MACS2 (version 2.1.1.20160309) was used to call peaks with default settings.

**Quantitative PCR Validation of the DNA-IP-Sequencing Experiment.** Immunoprecipitation of mitochondrial DNA using GFP-Trap (ChromoTek #gtma-20) was performed as described above. Four primer pairs encompassing the four peaks (target sites) identified by the DNA-IP-sequencing experiment, as well as six additional primer pairs amplifying sites that were not enriched (control sites) were designed as shown in *SI Appendix*, Table S2. SsoAdvanced Universal SYBR Green Supermix (BIO-RAD #1725270) was used for qPCR with a Realplex2 (Eppendorf) instrument. Each reaction was performed in triplicate (technical repeats) on two biological samples each for *Mito-GFP*, *35S::SHOT1-GFP* and *SHOT1p::SHOT1-GFP*. Percentage of input was calculated for each sample and normalized against *Mito-GFP* to obtain fold-enrichment values.

**Northern Blot Analysis of tRNAs.** 1 µg total RNA extracted from stage 1.02 seedlings (6) was separated on 1.3% agarose-formaldehyde gels and transferred to Hybond N^+^ nylon membrane (GE Healthcare). Membranes were UV-crosslinked at 120K microjoules/cm^2^, stained with 0.04% methylene blue in methanol for 10 min and destained with several changes of 25% ethanol. After staining, membranes were photographed and prehybridization was performed at 42 °C for 1 hour in Church buffer (7% SDS, 0.5 M NaPhosphate (pH 7.0), 1 mM EDTA). Fresh Church buffer with 20 nM of 5’ biotinylated oligo probes (Invitrogen, *SI Appendix*, Table S2) were added to the membranes and hybridized overnight at 42 °C. Membranes were washed once with 6X SSC, 0.1% SDS, twice with 4X SSC, 0.1% SDS and once with 2X SSD, 0.1% SDS at 42 °C for 15 min. each. Chemiluminescent nucleic acid detection module kit (Thermo Fisher Scientific) was used to detect the tRNAs.

**Immunoprecipitation of SHOT1-Interacting Proteins and Identification by Mass Spectrometry.** Purified mitochondria (0.5 or 1 mg mitochondrial proteins) from *Mito-GFP*, *35S::SHOT1-GFP*, and *SHOT1p::SHOT1-GFP* were resuspended in 400 µl IP buffer (20 mM Tris-HCl pH7.5, 150 mM NaCl, 1 mM EDTA, 0.5% Nonidet-P40, protease inhibitor cocktail (Sigma P9599)) and incubated for 30 min on ice. The lysate is clarified by centrifugation at 16000g for 10 min. The supernatant (~400 µl) was diluted with 600 µl of wash buffer (20 mM Tris-HCl pH7.5, 150 mM NaCl, 1 mM EDTA) to bring the concentration of NP40 to 0.2%. The lysate (1 ml) was added to 30 µl of GFP-Trap (ChromoTek #gta-20) equilibrated with the wash buffer and incubated with constant mixing at 4 °C for 30 min. The immunocomplexes on beads were washed 3 times with 1 ml wash buffer and eluted with 50 µl of 1X sample buffer by boiling for 10 min.

25 µl of the eluates were separated by SDS-PAGE for about 2 cm. The entire protein region of the gel was excised and subjected to in-gel trypsin digestion after reduction with DTT and alkylation with iodoacetamide. Peptides eluted from the gel were dried and resuspended in 10µL of 5% formic acid. The downstream analyses were performed at the mass spectrometry facility at University of Massachusetts Amherst. Samples were analyzed by LC-MS/MS using an Orbitrap Fusion mass spectrometer coupled to an Easy-nLC 1000 nano liquid chromatography system (Thermo Scientific), equipped with a nanoLC trap (75 um x 2 cm) and analytical column (PepMap RSLC C18 75 um x 15 cm). Mobile phases MPA: 0.1% formic acid in water, MPB: 0.1% formic acid in acetonitrile. Samples (5 ul) were loaded onto the trap column and desalted with 15 uL MPA, followed by transferring onto the analytical column and elution into the mass spectrometer with a gradient from 0-40% MPB over 90 min at a flow rate of 225 nl/min. MS spectra were acquired at a resolution setting of 60,000 (at 200 m/z) followed by data-dependent MS/MS spectra selecting top 10 most intense ions and fragmentation by CID with 35% normalized collision energy. Raw data were processed using MaxQuant version 1.5.5.1 (76) against the *Arabidopsis thaliana* (TAIR10) FASTA file (downloaded 06/2015). Search parameters included Trypsin/P specificity, up to 2 missed cleavages, a fixed modification of carbamidomethyl cysteine, and variable modifications of oxidized methionine, and N-terminal acetylation. Quantitative comparisons were made using spectral counts. Raw data were deposited to MassIVE repository (MSVxxxx).

Mitochondrial Nucleoid Staining, Confocal Microscopy, and Image Analysis. Mitochondrial DNA staining of plate-grown seedlings at stages 1.9-1.02 (6) was achieved by diluting PicoGreen solution (ThermoFisher Scientific) at 3 µl/ml directly into 0.5X MS medium and incubating for 45 min. Seedlings were then stained with the mitochondria selective dye MitoTracker Orange (ThermoFisher Scientific) by adding 500 nM directly to the medium and incubating for 10 min. Seedlings were washed in dye-free medium prior to observation under a confocal microscope (Olympus FluoView FV1000). PicoGreen signal was detected by excitation with a 473 nm laser and emission with a 490-540 nm band pass filter. MitoTracker Orange signal was detected by excitation with a 559 nm laser and emission with a 575-675 nm band pass filter. For image analysis, confocal images were opened with NIS-Elements package (Nikon). The areas of mitochondria and nucleoids were determined by General Analysis 3. The data were then exported and graphed using R studio.

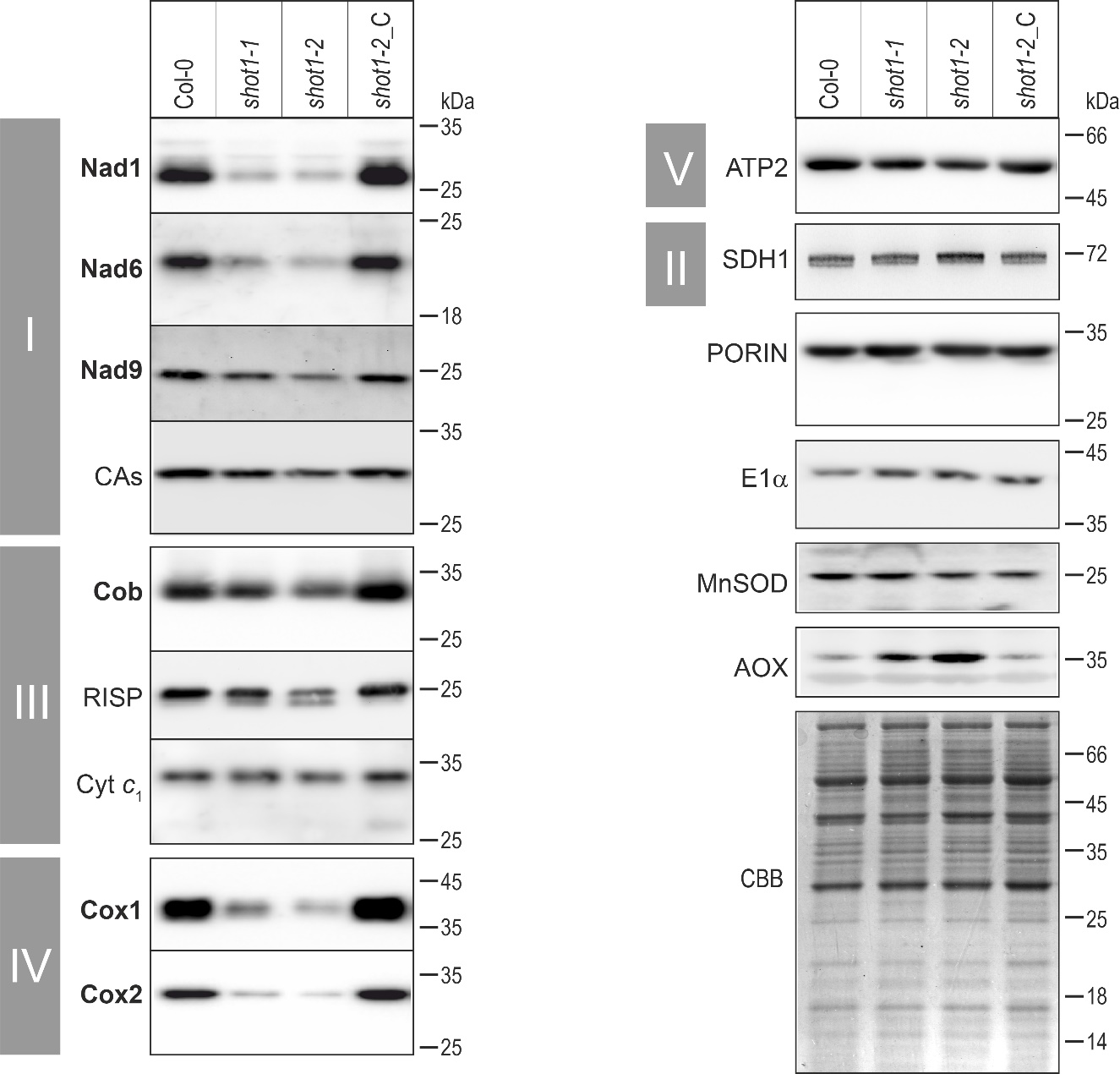

Fig. S1. Subunits of OXPHOS complexes containing mitochondria-encoded proteins are diminished in *shot1* mitochondria. Mitochondrial samples prepared from wild-type (Col-0), *shot1-1*, *shot1-2* plants, and from *shot1-2* complemented with the *SHOT1* wild-type allele (shot1-2_C) were resolved by SDS–PAGE and probed with antibodies against different mitochondrial proteins: Complex I subunits Nad1, Nad6, Nad9 and CA, complex II subunit SDH1, complex III subunits Cob, RISP and CYT *c_1_*, complex IV subunits Cox1 and Cox2, the complex V subunit ATP2, the outer membrane component PORIN, the E1α subunit of the pyruvate dehydrogenase complex, manganese superoxide dismutase (MnSOD), and alternative oxidase (AOX). Proteins written in bold are mitochondria-encoded; all other proteins are nucleus-encoded. The migration of size marker proteins is indicated on the right, the assignment of OXPHOS subunits to a specific complex is shown on the left. A Coomassie-stained gel (CBB) illustrates the comparability of mitochondrial samples prepared from the different genotypes.

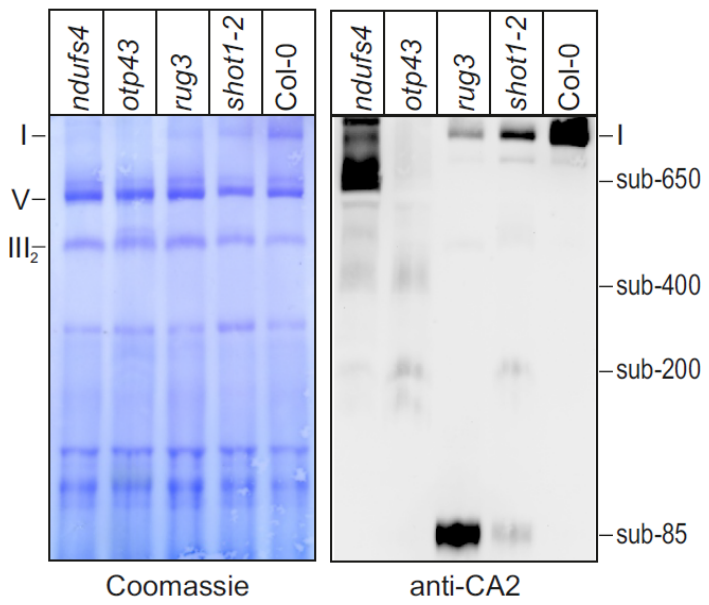

Fig. S2. *shot1-2* is defective in complex I assembly at the step of Nad2 insertion. To classify the major, low-molecular weight subcomplex over-accumulating in *shot1* mutants (see Fig. 1C), the subcomplex pattern was compared with wild-type plants (Col-0) and three different mutants impaired in the assembly of complex I: *ndufs4* over-accumulating the complex I membrane arm (sub-650) (20), *otp43* lacking the Nad1 subunit (21) and over-accumulating part of the membrane arm (sub-400, sub-200) (22), and *rug3* with diminished Nad2 expression and over-accumulation of the earliest detectable intermediate composed of the nucleus-encoded Carbonic Anhydrase (CA) and Carbonic Anhydrase-Like (CAL) subunits (23). Mitochondrial membrane complexes prepared from rosette-stage plants were resolved by BN-PAGE, transferred onto a PVDF membrane, and the membrane was probed with anti-CA2 antibodies, revealing assembly intermediates of the membrane arm (right panel). Comparable gel loading and protein transfer were controlled for by Coomassie-staining of the membrane (left panel). The smallest subcomplex over-accumulating in *shot1-2* is of the same size as the major subcomplex seen in *rug3*, showing that *shot1-2* is impaired in the insertion of Nad2 into complex I. Detection of sub-200 in *shot1-2* indicates additional defects at later steps of complex I assembly.

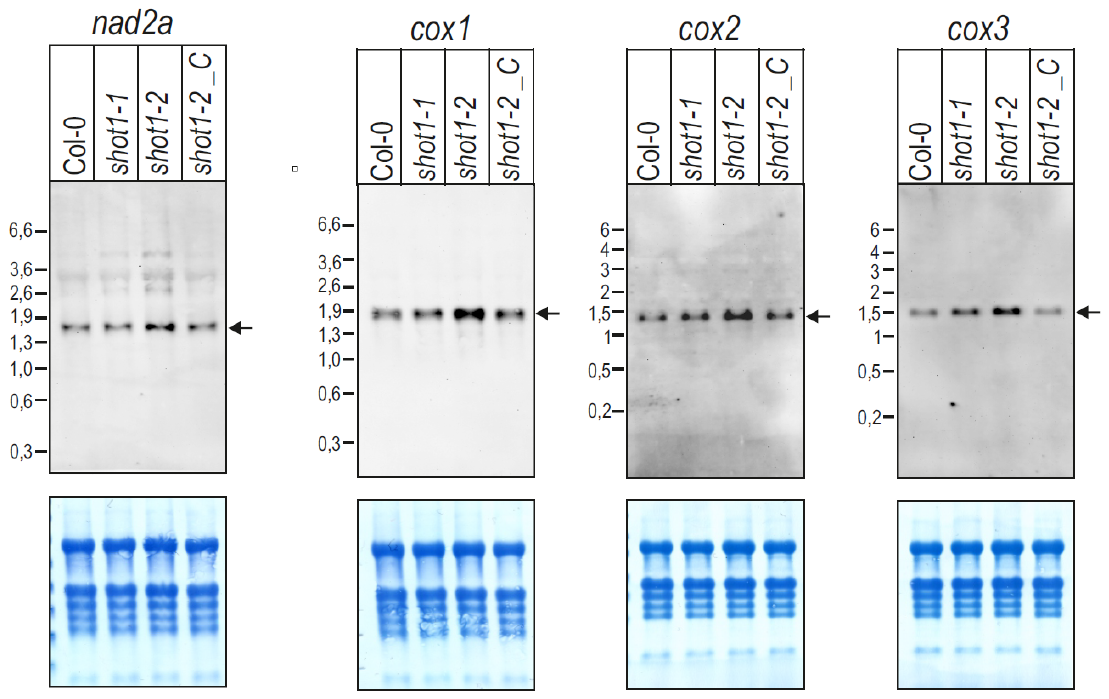

Fig. S3. SHOT1 does not affect the accumulation of correctly spliced or end-processed *nad2* and *cox* mRNAs. Probes for the mitochondrial *nad2, cox1, cox2* and *cox3* genes were hybridized to filter-immobilized total RNA isolated from wild-type (Col-0), *shot1-1*, *shot1-2* seedlings, and seedlings from *shot1-2* complemented with the *SHOT1* wild-type allele (shot1-2_C) (top panels). RNA marker sizes are indicated. Signals corresponding to the mature mRNAs are indicated by arrows. The same filters stained with methylene blue are shown as a loading control in the lower panels.

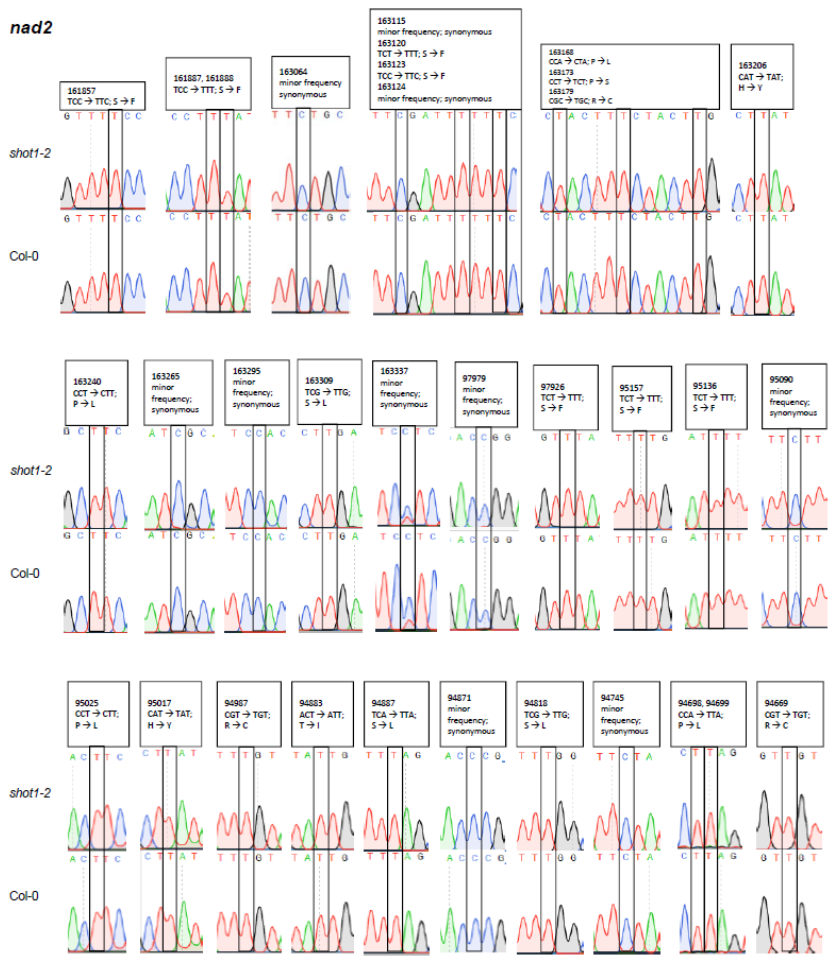

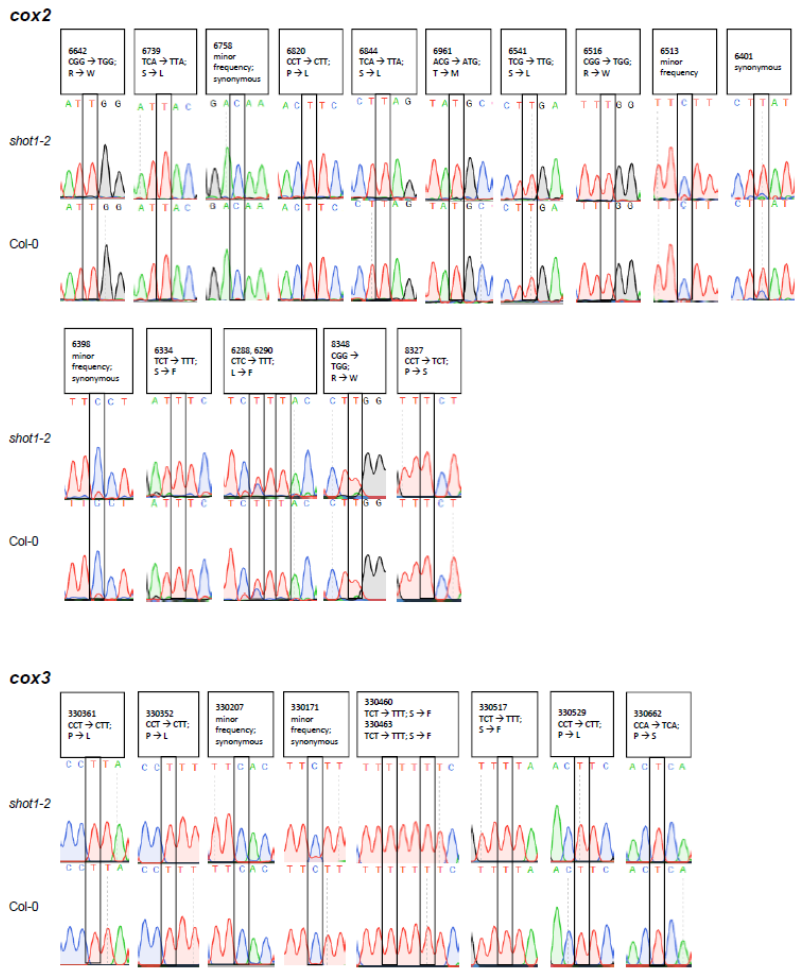

Fig. S4. *nad2* and *cox* mRNA editing is not impaired in *shot1-2*. mRNA editing was assessed through cDNA sequencing. Editing sites, as annotated in the *Arabidopsis thaliana* Col-0 genome (accession NC_037304.1) (24), are boxed in sequencing chromatograms. Numbers given above each editing sites refer to nucleotide positions in the Col-0 genome.

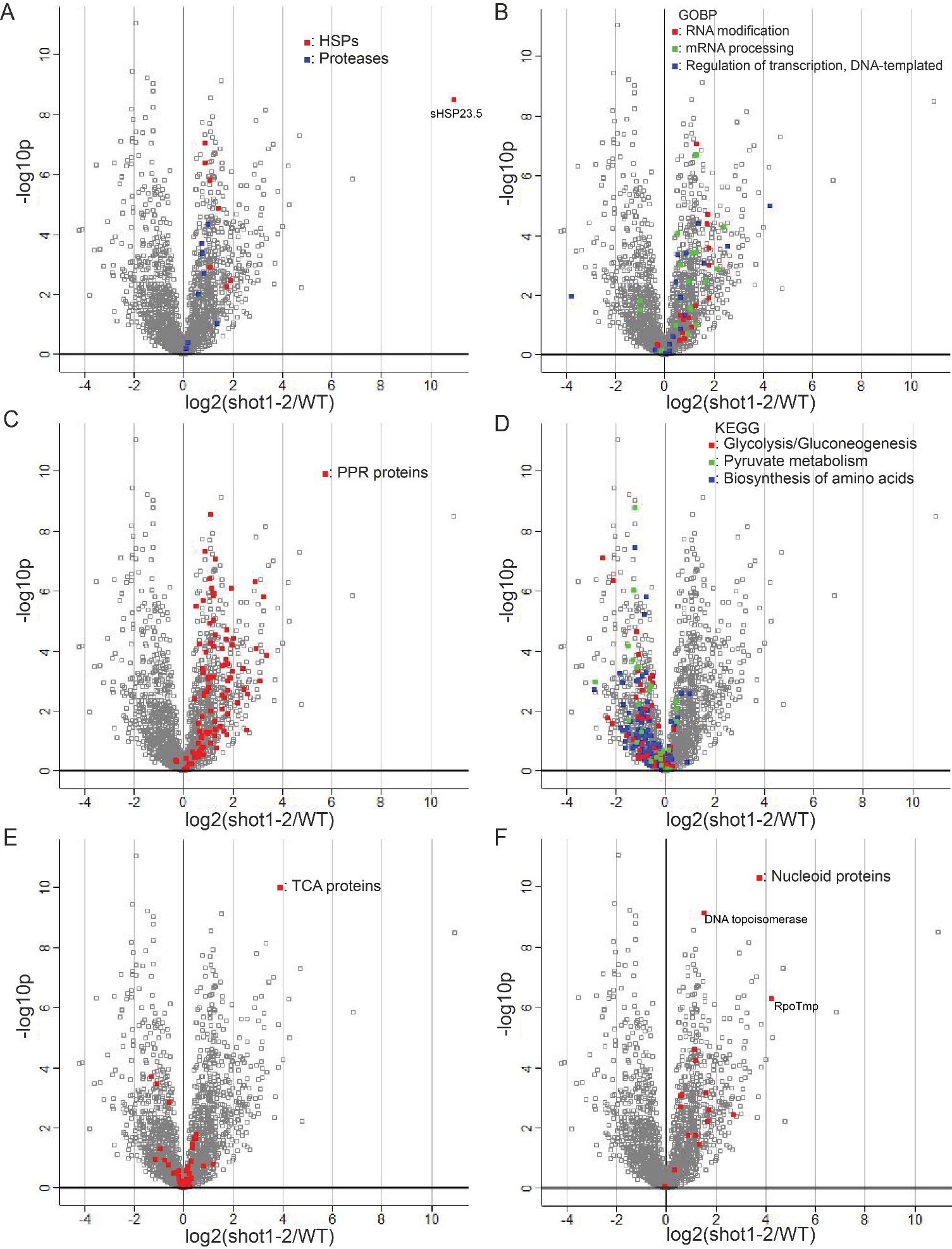

Fig. S5. Volcano plots of proteins in specific functional categories (GO terms and KEGG pathways). Each square represents a specific protein. Proteins in a specific category are colored as indicated in each plot. (A) Heat shock proteins (HSPs) and proteases are more abundant in the *shot1-2* mutant. (B) Proteins involved in transcription processes are increased in the *shot1-2* mutant. (C) PPR proteins are increased in the *shot1-2* mutant. (D) Proteins involved in carbon metabolism are decreased in the *shot1-2* mutant. (E) TCA proteins do not show a major difference in abundance between the wild type and *shot1-2* mutant. (F) Nucleoid proteins are increased in the *shot1-2* mutant. Please refer to Dataset S1 for identities of the proteins in volcano plots.

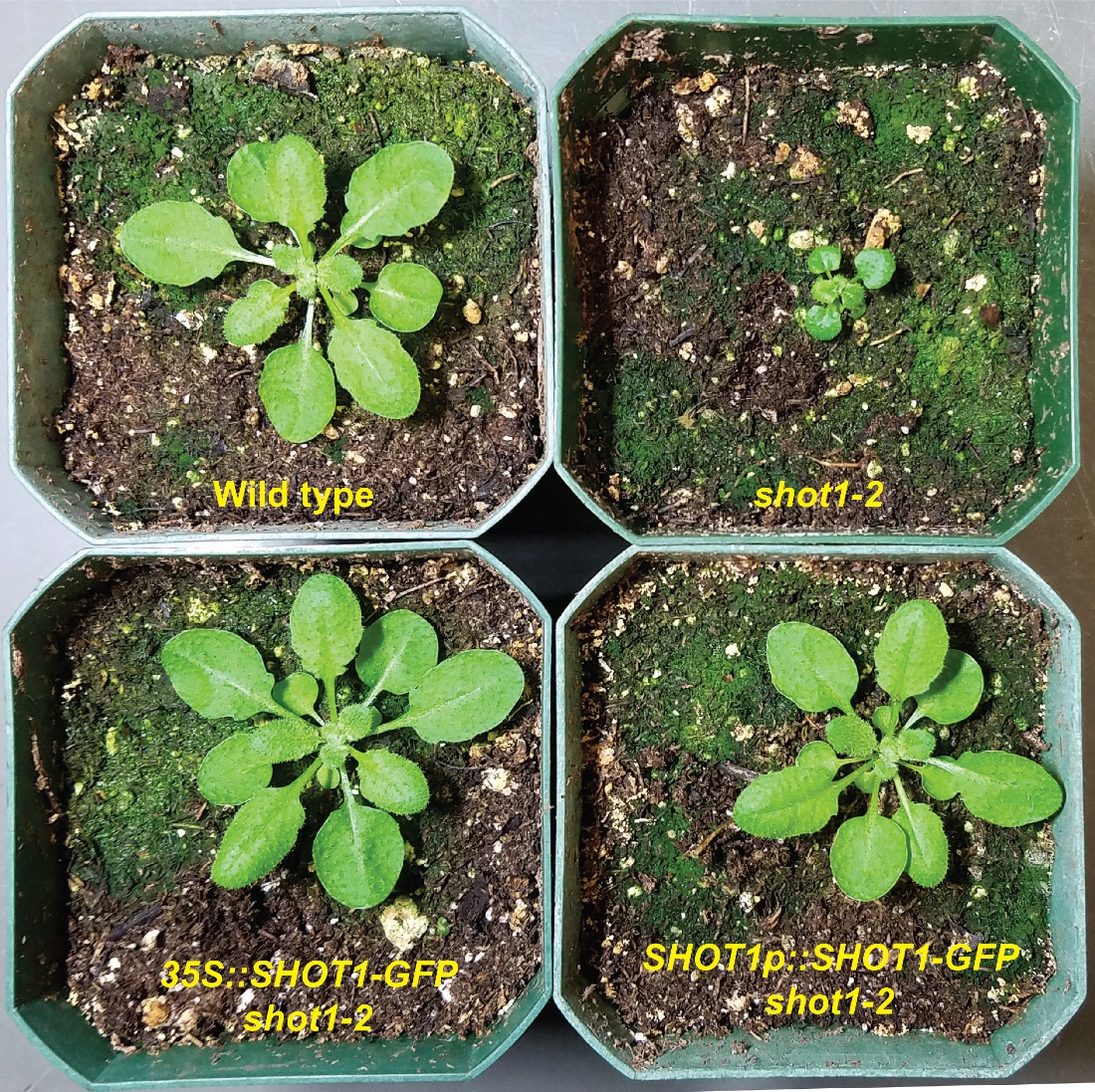

Fig. S6. Complementation of the *shot1-2* mutation by *35S::SHOT1-GFP* or *SHOT1p::SHOT1-GFP*. All plants are three weeks old.

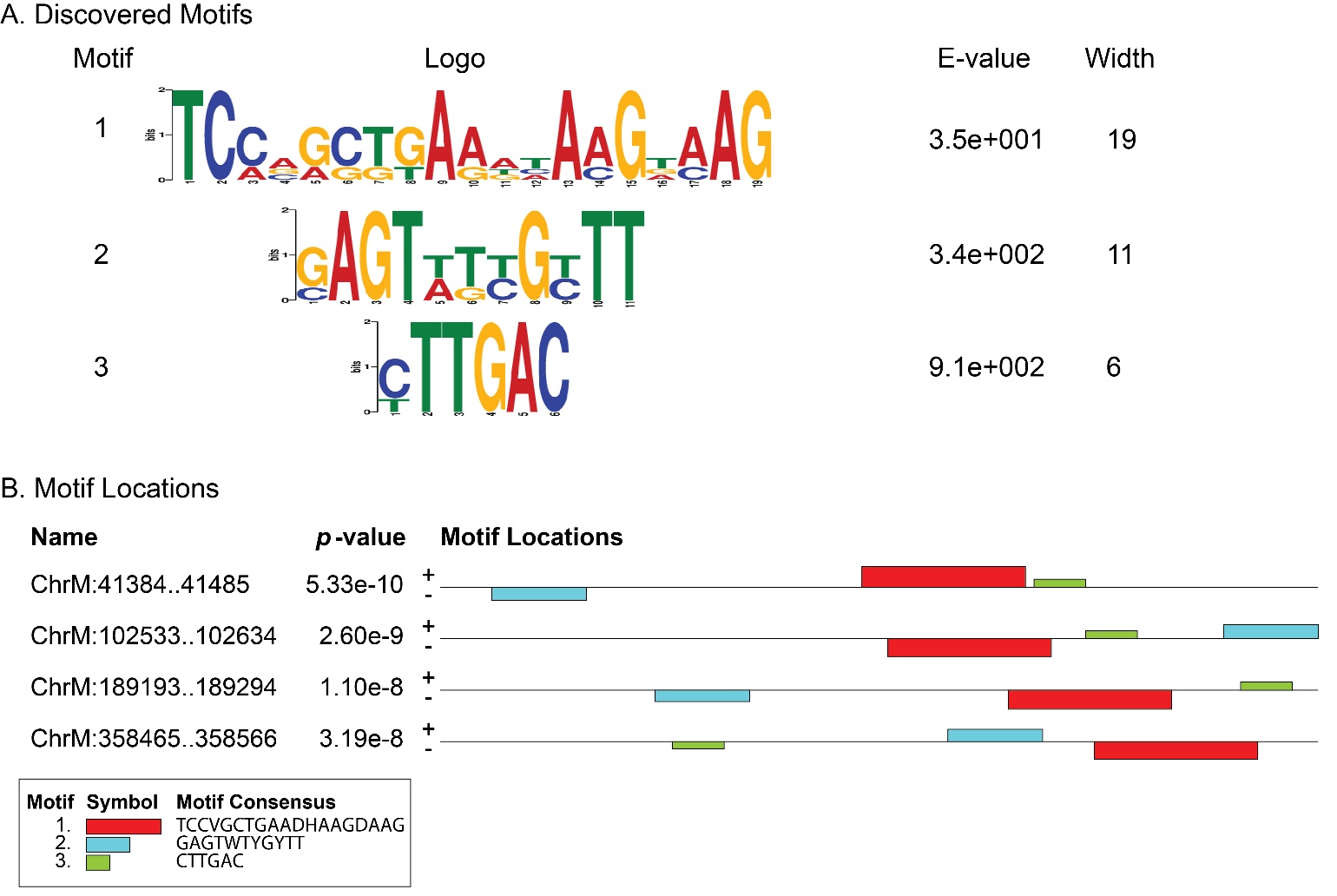

Fig. S7. Common motifs in the SHOT1 binding sites upstream of four tRNA genes. Conserved consensus sequences overrepresented in the 102-bp region surrounding the peaks of DNA-IP-Seq experiment were identified using the MEME server (25).

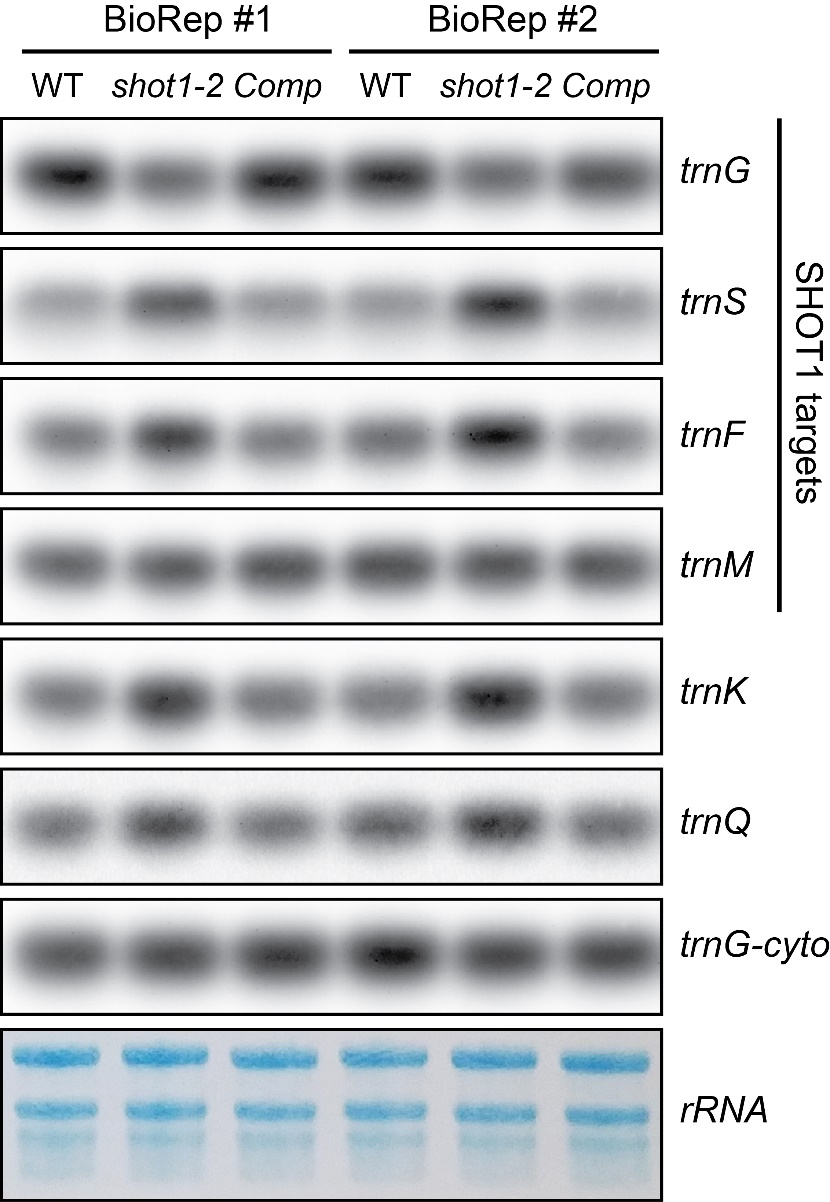

Fig. S8. RNA gel blot analysis of tRNA genes downstream of SHOT1 binding sites was performed with two biological replicate samples. Comp represents *shot1-2* complemented with the wild-type *SHOT1* gene. *trnK* and *trnQ* genes are included as a negative control because they are not related to SHOT1 binding sites. Cytosolic *trnG* (*trnG-cyto*) and methylene blue staining of ribosomal RNA are shown as a loading control.

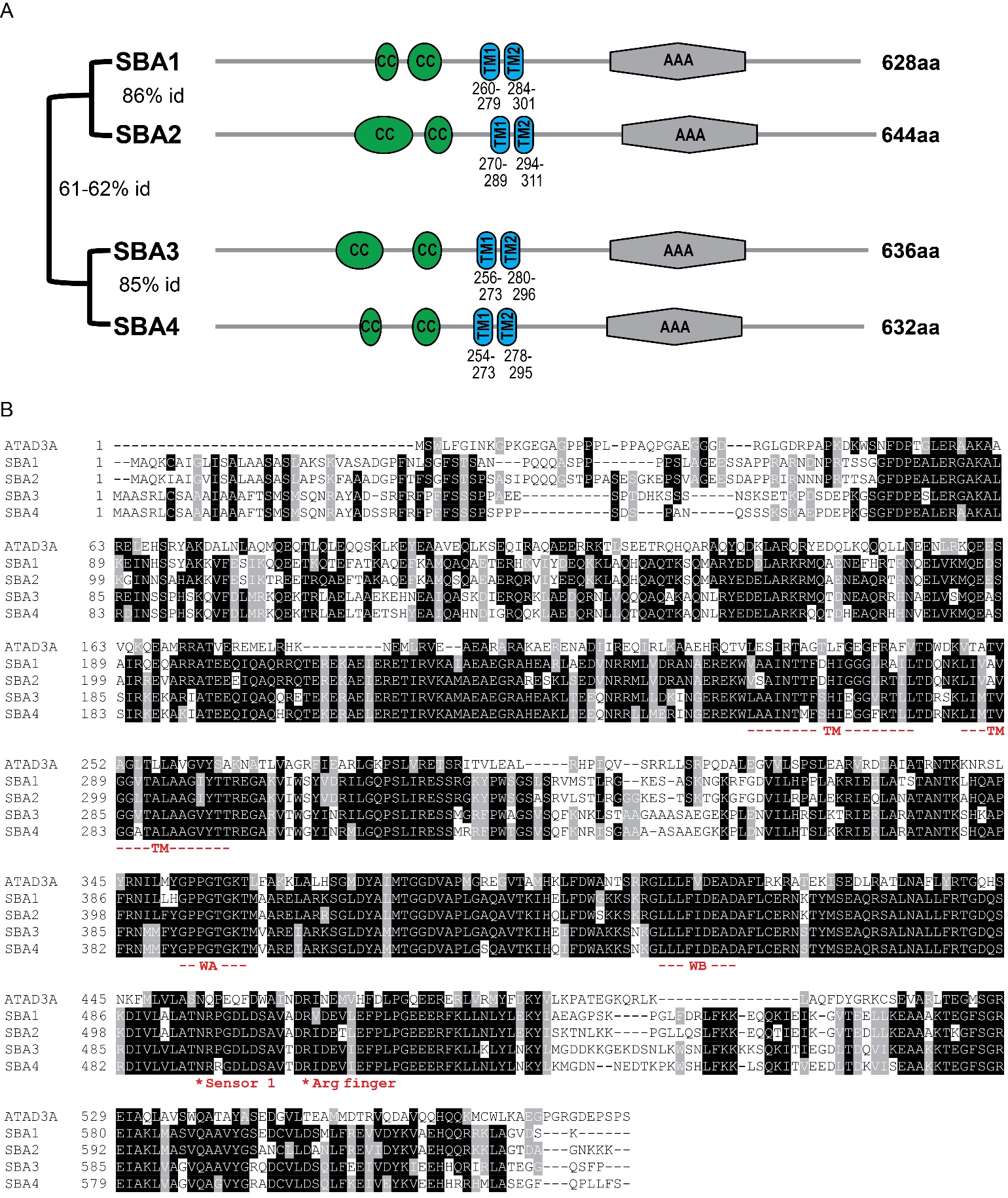

Fig. S9. Arabidopsis has four ATAD3 homologs, SBA1 (AT3G03060), SBA2 (AT5G16930), SBA3 (AT2G11330) and SBA4 (AT4G36580), in two clades that have a similar domain organization as animal ATAD3. (A) Domain organization and homology between SBA proteins. Protein homology is shown as a percentage of amino acid identity between the SBA proteins. Protein sequences were scanned with the InterPro website (https://www.ebi.ac.uk/interpro/) to find domains. The domain figures were created in MyDomains (https://prosite.expasy.org/mydomains/). CC: Coiled-Coil domains, TM: Transmembrane domains predicted by HMMTOP (26), AAA: ATPase Associated with diverse cellular Activities. (B) Protein sequence alignment of the human ATAD3A and SBA proteins was performed with Clustal Omega. BoxShade was used to make the image. WA: Walker A motif, WB: Walker B motif.

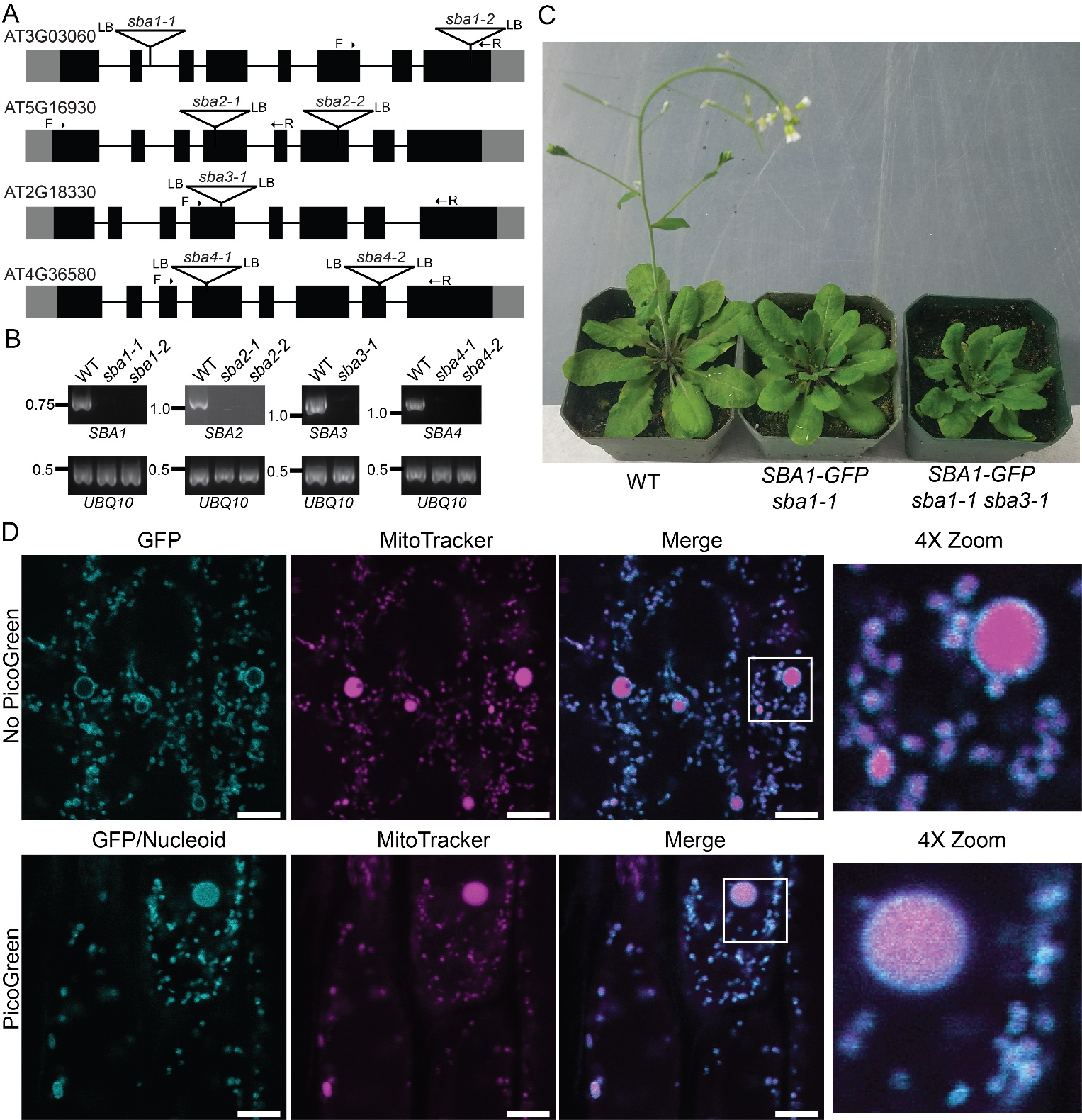

Fig. S10. *SBA1-GFP* plants show retarded growth and disturbed mitochondrial nucleoids. (A) Gene structures of *SBA* genes and positions of T-DNA insertions. Black boxes and gray boxes indicate coding and non-coding regions of mRNA, respectively. (B) RT-PCR analysis of the expression of SBA genes in T-DNA insertion mutants. RT-PCR was performed for 40 cycles using gene-specific primers whose locations are shown as arrows next to F (Forward) and R (Reverse) in (A). (C) 6-week-old *SBA1-GFP* plants in the *sba1-1* single or *sba1-1 sba3-1* double mutant background. (D) Root epidermal mitochondria of SBA1-GFP plants (*sba1-1* single mutant background) were observed without PicoGreen staining (upper panels) and with PicoGreen staining (lower panels). Boxed areas of each merged image are shown at higher magnification. Scale bars represent 5 µm.

**Table S1.** **SHOT1 binding sites were identified using MACS2 (version 2.1.1.20160309).** DNA-IP performed with 35S::SHOT1-GFP is shown.

| **start** | **end** | **length** | **abs_summit** | **pileup** | **-LOG10(pvalue)** | **fold_enrichment** | **-LOG10(qvalue)** | **Associated gene** |
| --- | --- | --- | --- | --- | --- | --- | --- | --- |
| 41299 | 41618 | 320 | 41435 | 37 | 13.27816 | 4.08129 | 10.23368 | trnS(GCT) |
| 102324 | 102816 | 493 | 102599 | 43 | 16.71946 | 4.52242 | 13.26443 | trnM(CAT) |
| 188987 | 189514 | 528 | 189240 | 49 | 22.36407 | 5.45774 | 16.79845 | trnG(GCC) |
| 358353 | 358658 | 306 | 358504 | 28 | 8.96821 | 3.49636 | 6.30474 | trnF |

Table S2. Oligonucleotides used in this study (5' to 3' direction).

| **Primers used for the synthesis of digoxigenin-labelled probes** | | |  |
| --- | --- | --- | --- |
| **Target** | **Primer 1** | **Primer 2** |  |
| *nad2* | CATTTTTTTATTGAGCCGAATCACT | TCCAAGCCAACCCACATTACTG |  |
| *cox1* | ATACCGAATCCAGGCAGAATGAG | GTAGGTAGCGGCACTGGGTG |  |
| *cox2* | CTGTCAAAAGTGAGTGACTGCTC | GGCAATTAGGATCTCAAGACGCAGC |  |
| *cox3* | AAGTTTGGGCCTCATATTTATCC | ATGATGAGCCCAAGTTACGG |  |
| *rrn18* | GCAAGTCGAACGTTGTTCTC | TACGCAGGCTCATCAAACAG |  |
| *rrn26* | TCAAAAGGCGAAAGTCTCGT | TTTTTCCTGGAAGTTTCAACC |  |
| **Primers used for the analysis of mRNA editing** | |  |  |
| **Target** | **PCR primer 1** | **PCR primer 2 (also used for RT)** | **Primers for Sanger sequencing of PCR products** |
| *cox1* | GGAGAATCAGGCAAGGTATGA | GTAGCTGCGGTGAAGTAGGC | GGAGAATCAGGCAAGGTATGA |
|  |  |  | ATCTGGTTCGATGGCTGTTC |
|  |  |  | TCTGGTGTTTCATCCATTTTAGG |
|  | TCTGGTGTTTCATCCATTTTAGG | TCGGCGACCTTTTCTTCTT | TCTGGTGTTTCATCCATTTTAGG |
|  |  |  | AGTTGGATCGCTACCATGTG |
| *cox2* | ACCAGCCATTTCCGTCTTC | CCCCTCCCTCACCTTACTCT | ATCCCGCAAAGGATTGTTC |
|  |  |  | GTGACTGCTCATCGGAACTG |
|  |  |  | CCCCTCCCTCACCTTACTCT |
| *cox3* | TCAATGCAATTAAAGAACCATCC | ATCAAAGGGAGTGGGAAAGC | TCAATGCAATTAAAGAACCATCC |
|  |  |  | ATCAAAGGGAGTGGGAAAGC |
| *nad2* | TGCTGGTCTAGCGATCCTTC | CCCTTTCTTTGAACCTTGTCA | TGCTGGTCTAGCGATCCTTC |
|  |  |  | CTACTCGCGGTATGCTCTTTATG |
|  |  |  | CGTGTTTCTATTTATGGTTCCTATG |
|  |  |  | TTCAGCATTACGGCAAACC |
| **Primers used for DNA IP-qPCR** | |  |  |
| **Target** | **Primer 1** | **Primer 2** |  |
| *ccmB* | GATCCGGAATGGATCGGTTAAA | TGCTGGATGTGATTCCAAGAG |  |
| *trnG* | GTCAAGAGCCGTGGTCCG | ACCATTAAGCTATTTCCGCCAG |  |
| *ccmFC* | CCAGACATAATTGGGAGAAGCTC | CCTTGGATTCGAGAAGACGG |  |
| *trnS* | AGCTATCAACTCAATTCTCTCGT | CTCCAAGTTGTTGATCGGAATTT |  |
| *trnF* | ACAGCATATTCCCAGTGAAGTAA | CTTCCTTACGGTACCCTTTCAG |  |
| *228k* | GAGATTCAGCTCTGATCACCAC | TCTAGGCCCAACGCATAAAC |  |
| *trnM* | GAAAAGAAGTCGCTCACCCG | ACCGCTGAGTCAAGTAGGTC |  |
| *trnM102k* | AGTACACTGAGTTAGGCGAGA | GGATAACCTACCTCTTCCGTGG |  |
| *trnS180k* | AGATTGTTTGTGGTTGTACCTTTAC | CCTACATTATGACCCAGAAACCT |  |
| *trnK* | AGACTGGAGCCTCGTAACA | GTCAATTAAGATTGGTGCGGAAA |  |
| **Sequences of 5' biotinylated probes used for tRNA blots** | |  |  |
| **Target** | **Sequence** |  |  |
| trnG | AGCGGAAGGAGGGACTTGAACCCTCA |  |  |
| trnS | CCAATGCCTTAAGCCACTCAGCCA |  |  |
| trnF | GATTCGAACCACTGCAAGGGTTTACAGTC |  |  |
| trnM | TCGCCGTATGAAAGCGATACTCTAACCG |  |  |
| trnK | TGGGTATAGCAGGACTTGAACCTGCG |  |  |
| trnQ | TGcATGCCGGTACCAAAAACCGG |  |  |
| trnG-cyto | TGCACCAGCCGGGAATCGAAC |  |  |
| **Primers used for genotyping** | |  |  |
| **Target** | **Primer 1** | **Primer 2** | **T-DNA specific primers** |
| sba1-1 | ATTCCTCTTACGCGAAAAAGG | TCTTGATACCCGAACATTCACC | CCCATTTGGACGTGAATGTAGACAC |
| sba1-2 | ATTCTTCTCCACGGTCCTCC | TTTTGAATCGACTCCAGCCAAT | CCCATTTGGACGTGAATGTAGACAC |
| sba2-1 | CCTACCAGAATCAACTTTGCAGC | AAGATGACTGAGTTCCTCAACCT | TAGCATCTGAATTTCATAACCAATCTCGATACAC |
| sba2-2 | TGTGGCAGTTTACCCTGGTAG | GGAAGAGGAGGGCATTTAGTG | ATTTTGCCGATTTCGGAAC |
| sba3-1 | GTCTGGGAAGCGACAAATATGA | AGGCTTCTTAAATCACAGCTGT | ATTTTGCCGATTTCGGAAC |
| sba4-1 | ATGGGGCATTTGAAATGGGC | GCCCCTGAGATCCTGTTCTT | ATTTTGCCGATTTCGGAAC |
| sba4-2 | GGTATGCATCTCCGAACCAAA | TGGAAGTGGGAACTCAATGAC | ATTTTGCCGATTTCGGAAC |
| **Primers used for RT-PCR** | |  |  |
| **Target** | **Primer 1** | **Primer 2** |  |
| UBQ10, AT4G05320 | GATCTTTGCCGGAAAACAATTGGAGGATGGT | CGACTTGTCATTAGAAAGAAAGAGATAACAGG |  |
| SBA1 | ATTCTTCTCCACGGTCCTCC | TTTTGAATCGACTCCAGCCAAT |  |
| SBA2 | ATGGCTCAGAAAATTGCGATTG | TGATCTGTTAGAATGGTACGCAAC |  |
| SBA3 | GATAATGAAGCTCAGAGACGGC | ACCTTTCTTGTCGTCACCCATT |  |
| SBA4 | GAGGACCAGAGAAATCTTTTGCA | TGGAAGTGGGAACTCAATGAC |  |

**Dataset S1**. **Mitochondrial proteomics of *shot1-2*.** Proteins shown in the volcano plots of Fig. 3 and Fig. S5 are listed.
